# Loss of Tsc1 from striatal direct pathway neurons impairs endocannabinoid-LTD and enhances motor routine learning

**DOI:** 10.1101/2019.12.15.877126

**Authors:** Katelyn N. Benthall, Katherine R. Cording, Alexander H.C.W. Agopyan-Miu, Emily Y. Chen, Helen S. Bateup

## Abstract

Tuberous Sclerosis Complex (TSC) is a neurodevelopmental disorder in which patients frequently present with autism spectrum disorder (ASD). A core diagnostic criterion for ASD is the presence of restricted, repetitive behaviors, which may result from abnormal activity in striatal circuits that mediate motor learning, action selection and habit formation. Striatal control over motor behavior relies on the coordinated activity of two subtypes of principle neurons, direct pathway and indirect pathway spiny projection neurons (dSPNs or iSPNs, respectively), which provide the main output of the striatum. To test if altered striatal activity is sufficient to cause changes to motor behavior in the context of TSC, we conditionally deleted *Tsc1* from dSPNs or iSPNs in mice and determined the consequences on synaptic function and motor learning. We find that mice with loss of Tsc1 from dSPNs, but not iSPNs, have enhanced motor routine learning in the accelerating rotarod task. In addition, dSPN Tsc1 KO mice have impaired endocannabinoid-mediated long-term depression (eCB-LTD) at cortico-dSPN synapses in the dorsal striatum. Consistent with a loss of eCB-LTD, disruption of Tsc1 in dSPNs, but not iSPNs, results in a strong enhancement of corticostriatal synaptic drive. Together these findings demonstrate that within the striatum, dSPNs show selective sensitivity to Tsc1 loss and indicate that enhanced cortical activation of the striatal direct pathway is a potential contributor to altered motor behaviors in TSC.

## Introduction

Autism spectrum disorder (ASD) is characterized by social and communication deficits, as well as the presence of restricted and repetitive behaviors (RRBs). Over a hundred genes have been associated with ASD risk (Satterstrom et al., 2020), including genes that cause syndromic disorders in which ASD is frequently diagnosed, together with a constellation of other neurological, psychiatric and medical conditions (Sztainberg and Zoghbi, 2016). One such syndrome is Tuberous Sclerosis Complex (TSC), a disorder caused by mutations in either *TSC1* or *TSC2*, in which about half of patients meet the clinical criteria for an ASD diagnosis and most have epilepsy and other behavioral or cognitive disorders (Curatolo et al., 2015; Davis et al., 2015). The TSC1 and TSC2 proteins form a complex that negatively regulates the mTORC1 signaling pathway, a central signaling hub controlling cellular metabolic processes such as protein and lipid synthesis and autophagy (Saxton and Sabatini, 2017). When the TSC1/2 complex is disrupted, mTORC1 signaling is constitutively active, leading to excessive cell growth and altered cellular metabolism (Huang and Manning, 2008). Dysregulation of mTORC1 signaling is not limited to TSC but may occur commonly in ASD (de Vries, 2010; Kelleher and Bear, 2008; Tang et al., 2014; Winden et al., 2018).

While epilepsy in TSC likely arises from altered excitability in forebrain circuits, the brain regions and cell types important for ASD-related behaviors in individuals with TSC are less well understood. We hypothesized that alterations in striatal circuits, which mediate motor learning, action selection, and habit formation, might contribute to the RRBs observed in TSC patients with ASD. Indeed, striatal alterations have been implicated in non-syndromic forms of ASD by structural and functional MRI studies (Di Martino et al., 2011; Qiu et al., 2010; Turner et al., 2006). Further, in a study of TSC children with and without ASD, striatal metabolism was found to differ specifically in those children with ASD (Asano et al., 2001). Notably, this altered striatal activity was correlated with the presence of RRBs in the TSC children with ASD (Asano et al., 2001). Recently, the presence of self-injurious behavior in individuals with TSC was shown to be associated with structural changes in basal ganglia nuclei (Gipson et al., 2019).

Mouse studies support the link between striatal alterations and ASD-related behaviors (Fuccillo, 2016; Li and Pozzo-Miller, 2019; Rothwell, 2016), showing that mutations in ASD-risk genes can endow the striatum with an enhanced ability to acquire fixed motor routines (Kwon et al., 2006; Nakatani et al., 2009; Penagarikano et al., 2011; Platt et al., 2017; Rothwell et al., 2014). Changes in striatal synaptic properties have been reported in multiple mouse models with mutations in ASD-risk genes (Fuccillo, 2016; Li and Pozzo-Miller, 2019; Peca et al., 2011; Peixoto et al., 2016). Our group has previously shown input- and cell type-specific changes in synaptic transmission in striatal neurons with postnatal loss of Tsc1 (Benthall et al., 2018). Therefore, the goal of this study was to determine whether deletion of *Tsc1* selectively in striatal neurons is sufficient to alter motor behaviors relevant to ASD and TSC.

Striatal function depends on the coordinated activity of two subpopulations of GABAergic principal cells: direct pathway spiny projection neurons (dSPNs) and indirect pathway spiny projection neurons (iSPNs). dSPNs preferentially express D1 dopamine receptors, while iSPNs express D2 dopamine receptors and A2a adenosine receptors (Gerfen and Surmeier, 2011). dSPNs and iSPNs are distinct in their projection targets, diverging as they leave the striatum to innervate the globus pallidus internal segment (GPi) and substantia nigra pars reticulata (SNr), or the globus pallidus external segment (GPe), respectively (Gerfen and Surmeier, 2011). At the simplest level, activation of dSPNs increases motor behaviors and action selection while stimulation of iSPNs inhibits movement (Bateup et al., 2010; Kravitz et al., 2010; Tai et al., 2012). Therefore, coordinated activity between dSPNs and iSPNs allows for the selection of appropriate actions to be performed for any given context, while inappropriate actions are suppressed. Cortical and thalamic inputs make the primary glutamatergic synapses onto SPNs (Ding et al., 2008; Doig et al., 2010), and modification of the strength of this excitatory drive by long-term synaptic plasticity mediates motor learning and habit formation (Graybiel and Grafton, 2015; Gremel and Lovinger, 2017).

To determine how developmental loss of Tsc1 affects striatal synapses and whether this is sufficient to alter striatal-dependent behaviors, we generated mice in which *Tsc1* was selectively deleted from either dSPNs or iSPNs. We found that motor routine learning was enhanced in mice with *Tsc1* deletion restricted to dorsal striatal dSPNs, but not iSPNs, which also occurred in mice with global haploinsufficiency of *Tsc2*. Further, we found that loss of Tsc1 caused a non-cell autonomous enhancement in corticostriatal drive of dSPNs but not iSPNs, which was associated with loss of endocannabinoid-mediated long-term depression (eCB-LTD). Together, these findings demonstrate that increased corticostriatal drive of dSPNs resulting from Tsc1 loss is sufficient to enhance the establishment of fixed motor routines.

## Results

### Upregulation of mTORC1 and somatic hypertrophy in SPNs with *Tsc1* deletion

To test whether striatal-specific disruption of the Tsc1/2 complex is sufficient to alter motor behaviors, we bred conditional *Tsc1* knock-out mice (*Tsc1^fl/fl^*) (Kwiatkowski et al., 2002) with striatal-restricted *Drd1-Cre* or *Adora2a-Cre* mice (Gong et al., 2007) to achieve dSPN- or iSPN-specific loss of Tsc1, respectively (Fig. 1a,b). In addition, we bred in a Cre-dependent tdTomato fluorescent reporter (Ai9) to visualize Cre-expressing cells (Madisen et al., 2010). We used the EY217 *Drd1-Cre* founder line from GENSAT as it has more striatal-restricted expression than the commonly used EY262 line (Fig. S1a,b). In preliminary studies, we found that *Tsc1^fl/fl^;Drd1-Cre(EY262)* mice died prematurely, around postnatal day 15, which was not the case for Cre-positive *Tsc1^wt/wt^* or *Tsc1^wt/fl^* animals. The premature mortality may have been caused by seizures due to cortical Cre expression in this line (Fig. S1b), as loss of Tsc1 from only a small percent of cortical neurons is sufficient to induce seizures (Lim et al., 2017). *Tsc1^fl/fl^;Drd1-Cre(EY217)* did not exhibit premature mortality, likely due to the more striatal-restricted Cre expression. Notably, Cre expression in *Drd1-Cre(EY217)* mice was highest in the dorsal striatum, with 40-45% of neurons showing Cre-dependent recombination, whereas less than 8% of neurons in the ventral striatum exhibited Cre recombinase activity (Fig. S1c-f, k). *Adora2a-Cre* was expressed uniformly throughout the striatum in approximately 45% of neurons (Fig. S1g-j, l). Homozygous loss of *Tsc1* did not affect the number of tdTomato+ dSPNs or iSPNs across striatal regions (Fig. S1k,l).

**Figure 1.**
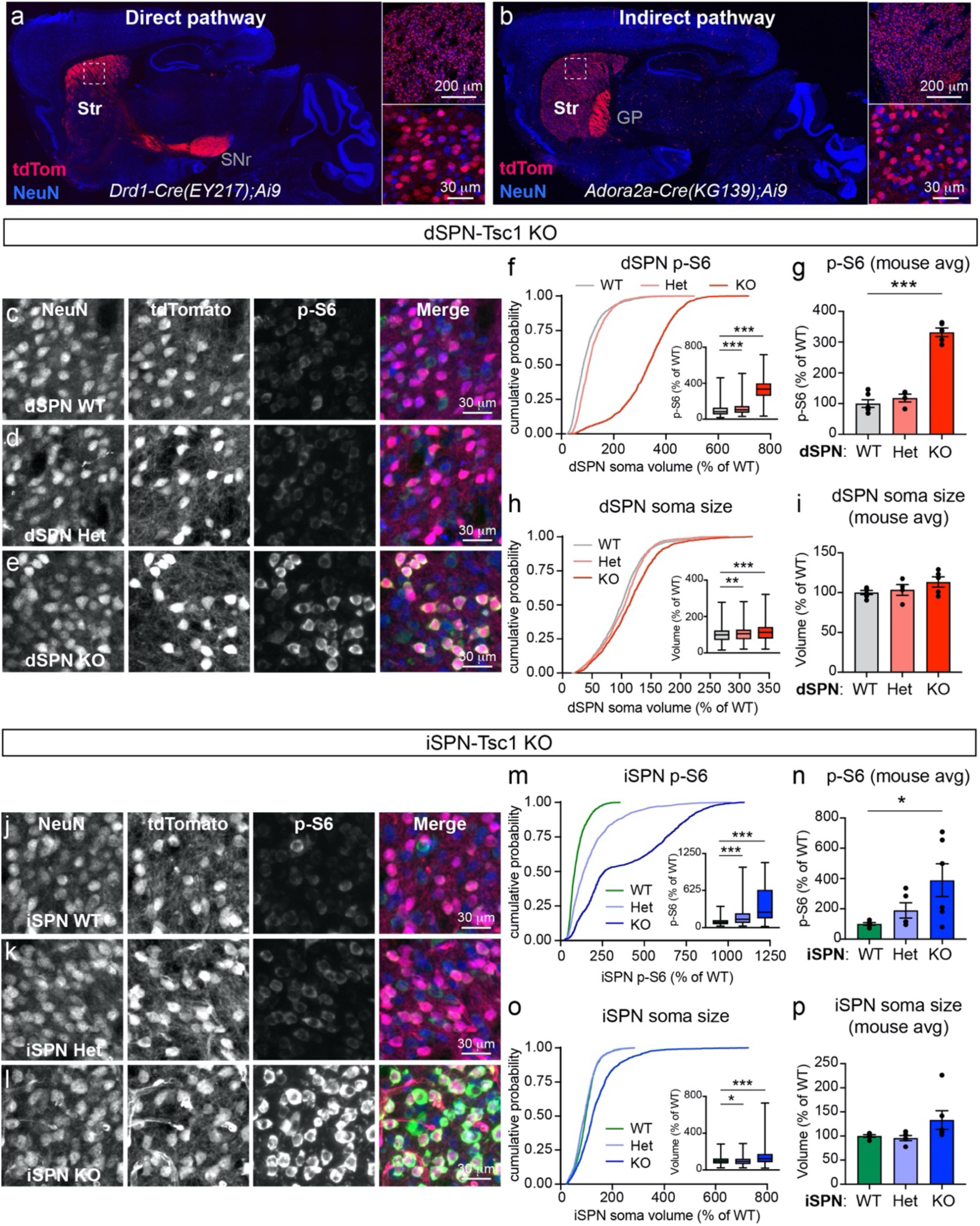
Developmental loss of *Tsc1* from striatal projection neurons induces mTORC1 activation. (a,b) Images of sagittal brain sections showing the expression pattern of a tdTomato Cre reporter in a *Drd1-Cre;Ai9* mouse (EY217 GENSAT founder line, a) and an *Adora2a-Cre;Ai9* mouse (b) with selective expression in the direct and indirect pathways, respectively. Right panels show higher magnification images of the dorsal striatum (boxed region). NeuN antibody staining is in blue. Str=striatum, SNr=substantia nigra pars reticulata, GP=globus pallidus (c-e) Confocal images of dorsolateral striatum from a dSPN-Tsc1 WT (c), Tsc1 Het (d), and Tsc1 KO (e) mouse labeled with antibodies against phosphorylated S6 (p-S6, Ser240/244, green in the merged image) and NeuN (blue in the merged image). TdTomato (red in the merged imaged) is expressed in dSPNs. (f) Cumulative distribution of dSPN p-S6 fluorescence intensity per cell, expressed as a percentage of wild-type (WT). 1800 cells from 6 dSPN-Tsc1 WT mice, 1200 cells from 4 dSPN-Tsc1 Het mice, and 1500 cells from 5 dSPN-Tsc1 KO mice were analyzed. Inset box plots display the 25-75%-ile p-S6 per cell by genotype (line at the median, whiskers = min to max). Kruskal-Wallis, p<0.0001; Dunn’s multiple comparisons tests, WT vs Het, ***, p<0.0001; WT vs KO, ***, p<0.0001. (g) Bar graphs display the mean +/− SEM p-S6 level per mouse for each genotype normalized to WT (calculated from the data in f). Dots represent individual mice. One-way ANOVA, p<0.0001; Holm-Sidak’s multiple comparisons tests, WT vs Het, p=0.3701; WT vs KO, ***, p<0.0001. (h) Cumulative distribution of dSPN soma volume per cell, measured from the same cells as in panel f, expressed as a percentage of wild-type (WT). Inset box plots display the 25-75%-ile soma volume per cell by genotype (line at the median, whiskers = min to max). Kruskal-Wallis, p<0.0001; Dunn’s multiple comparisons tests, WT vs Het, **, p=0.0071; WT vs KO, ***, p<0.0001. (i) Bar graphs display the mean +/− SEM soma volume per mouse for each genotype normalized to WT (calculated from the data in h). Dots represent individual mice. One-way ANOVA, p=0.1927; Holm-Sidak’s multiple comparisons tests, WT vs Het, p=0.6639; WT vs KO, p=0.1557. (j-l) Confocal images of dorsolateral striatum from an iSPN-Tsc1 WT (j), Tsc1 Het (k), and Tsc1 KO (l) mouse labeled with antibodies against phosphorylated S6 (p-S6, Ser240/244, green in the merged image) and NeuN (blue in the merged image). TdTomato (red in the merged imaged) is expressed in iSPNs. (m) Cumulative distribution of iSPN p-S6 fluorescence intensity per cell, expressed as a percentage of wild-type (WT). 1500 cells from 5 iSPN-Tsc1 WT mice, 1500 cells from 5 iSPN-Tsc1 Het mice, and 1800 cells from 6 iSPN-Tsc1 KO mice were analyzed. Inset box plots display the 25-75%-ile p-S6 per cell by genotype (line at the median, whiskers = min to max). Kruskal-Wallis, p<0.0001; Dunn’s multiple comparisons tests, WT vs Het, ***, p<0.0001; WT vs KO, ***, p<0.0001. (n) Bar graphs display the mean +/− SEM p-S6 level per mouse for each genotype normalized to WT (calculated from the data in m). Dots represent individual mice. One-way ANOVA, p=0.0481; Holm-Sidak’s multiple comparisons tests, WT vs Het, p=0.4393; WT vs KO, *, p=0.0368. (o) Cumulative distribution of iSPN soma volume per cell, measured from the same cells as in panel m, expressed as a percentage of wild-type (WT). Inset box plots display the 25-75%-ile soma volume per cell by genotype (line at the median, whiskers = min to max). Kruskal-Wallis, p<0.0001; Dunn’s multiple comparisons tests, WT vs Het, *, p=0.0142; WT vs KO, ***, p<0.0001. (p) Bar graphs display the mean +/− SEM soma volume per mouse for each genotype normalized to WT (calculated from the data in o). Dots represent individual mice. Kruskal-Wallis, p=0.0242; Dunn’s multiple comparisons tests, WT vs Het, p>0.9999; WT vs KO, p=0.0687.

To confirm that *Tsc1* deletion upregulated mTORC1 signaling in SPNs, we quantified phosphorylation of the mTORC1 pathway target ribosomal protein S6 in dorsal striatal neurons from *Tsc1^fl/fl^;Drd1-Cre;Ai9* (referred to as dSPN-Tsc1 KO) and *Tsc1^fl.fl^;Adora2a-Cre;Ai9* (referred to as iSPN-Tsc1 KO) mice. Loss of Tsc1 caused a gene dose-sensitive increase in p-S6 levels in both dSPNs and iSPNs, indicating mTORC1 pathway hyperactivity in both cell types (Fig. 1c-g and j-n). Prior studies have demonstrated pronounced somatic hypertrophy in Tsc1 KO neurons in various brain regions, consistent with the known role of mTORC1 in regulating cell size (Bateup et al., 2011; Feliciano et al., 2011; Kosillo et al., 2019; Malik et al., 2019; Normand et al., 2013; Tsai et al., 2012). In the striatum, we observed a relatively modest but significant increase in the soma volume of dSPN-Tsc1 KO and iSPN-Tsc1 KO neurons (Fig. 1h,i and o,p). Analysis of the distribution of dSPN-Tsc1 Het cell sizes revealed a small increase in soma volume compared to wild-type (WT) cells (Fig. 1h), while iSPN-Tsc1 Het cells had slightly reduced soma volume (Fig. 1o). Together these results show that complete loss of Tsc1 from SPNs results in robust activation of mTORC1 signaling leading to moderate somatic hypertrophy. Heterozygous loss of Tsc1 mildly increases mTORC1 activity in dSPNs and iSPNs but does not strongly affect soma size.

### Loss of Tsc1 from dSPNs but not iSPNs enhances motor routine learning

To determine whether SPN-specific loss of Tsc1 was sufficient to alter motor behaviors we investigated general locomotor activity and self-grooming behavior in the open field. We found no significant differences in the total distance traveled, number of rears, or grooming bouts in dSPN- or iSPN-Tsc1 Het or KO mice compared to controls (Fig. S2a-f). These results suggest that loss of Tsc1 from a single SPN subtype is not sufficient to alter gross motor behavior or induce spontaneous stereotypies. A summary of the behavior test results by genotype and sex are shown in Supplemental Table 1.

The accelerating rotarod task is a striatal-dependent motor learning assay (Yin et al., 2009) that is altered in multiple mouse models with mutations in ASD-risk genes (Fuccillo, 2016). In this test, mice learn to run on a rod revolving at increasing speed over four days of training with three trials performed each day (Rothwell et al., 2014). Over the course of training, mice develop a stereotyped motor routine to stay on the apparatus for increasing amounts of time and thus reach higher terminal speeds. Control mice improve at the task over training, with an average learning rate of 2-3 rpm/trial. We found that dSPN-Tsc1 Het and KO mice had similar initial rotarod performance as WT littermates on the first trial, reflecting normal baseline motor coordination (Fig. 2a,b). However, dSPN-Tsc1 KO mice exhibited significantly enhanced motor learning, measured by the slope of performance from the first to last trial for each mouse, compared to littermate controls (Fig. 2a,c). dSPN-Tsc1 Het animals also displayed a mild enhancement of motor learning consistent with a gene dose-dependent effect (Fig. 2a). The enhancement in rotarod performance was most pronounced for the more challenging 10-80 rpm acceleration trials. Across these trials (trials #7-12), 72% and 70% of dSPN-Tsc1 Het and KO mice, respectively, displayed continued improvement indicated by a positive slope of their learning curve (Fig. 2d). This contrasted with the less than 50% of dSPN-Tsc1 WT mice that showed improvement across trials 7-12, reflecting a close to ceiling level of performance by trial 7 (Fig. 2d). These results are unlikely to be driven by changes in weight as weight was not significantly different between genotypes, although dSPN-Tsc1 KO mice tended to be lighter (dSPN-Tsc1 WT male = 34.55 +/− 2.42g, Het male = 33.11 +/− 1.65g, KO male = 29.20 +/− 1.35g, p=0.3453, Kruskal-Wallis test; dSPN-Tsc1 WT female = 31.30 +/− 1.99g, Het female = 26.50 +/− 1.22g, KO female = 25.44 +/− 1.77g; p=0.0910, Kruskal-Wallis test). Further, we found that weight was not significantly correlated with rotarod learning rate for either female or male mice (Fig. S3). We examined rotarod performance in mice with loss of Tsc1 in iSPNs and strikingly, we found no difference in either initial motor performance, motor learning ability, or performance on the 10-80 rpm trials between genotypes (Fig. 2e-h).

**Figure 2.**
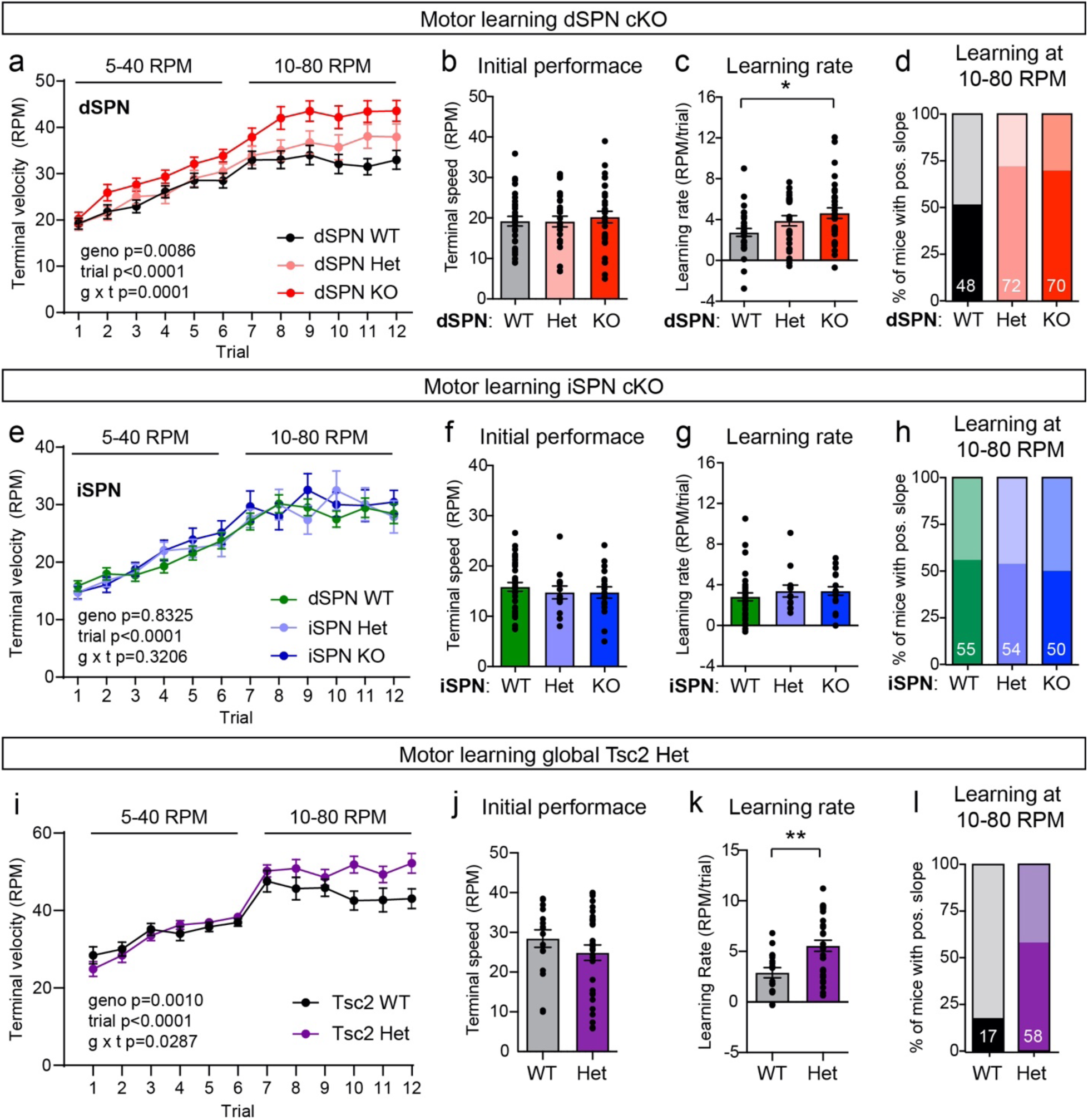
Loss of Tsc1 from dSPNs enhances motor routine learning. (a) Performance on the accelerating rotarod across 12 trials (4 days) in dSPN-Tsc1 WT (n=31), Het (n=26), and KO (n=33) mice. Circles represent mean +/− SEM. Mixed-effects analysis p values are shown. RPM = revolutions per minute. (b) Rotarod performance on trial 1 for dSPN-Tsc1 WT, Het and KO mice quantified as terminal speed. Bars represent mean +/− SEM. Dots represent individual mice (n is the same as in panel a). Kruskal-Wallis test, p=0.7794. (c) Overall learning rate of dSPN-Tsc1 WT, Het and KO mice calculated as the slope of the line of performance on the first trial (1) to the last trial (12) for each mouse (RPM/trial). Bars represent mean +/− SEM. Dots represent individual mice (n is the same as in panel a). Kruskal-Wallis test, p=0.0343; Dunn’s multiple comparisons tests, dSPN WT vs KO, *, p=0.0206; dSPN WT vs Het, p=0.2199. (d) Percentage of dSPN-Tsc1 WT, Het, and KO mice with a positive learning curve (slope of performance) from trial 7 to trial 12 (10-80 RPM acceleration trials). Dark shaded regions indicate the percent of mice with a positive slope, light shaded regions indicate the percent of mice with a negative slope, within each genotype. The numbers on the bars indicate the percentage of mice of each genotype with a positive slope. N is the same as for panel a. (e) Performance on the accelerating rotarod across 12 trials (4 days) in iSPN-Tsc1 WT (n=34 mice), Het (n=13 mice), and KO (n=18 mice). Dots represent mean +/− SEM. Two-way ANOVA p values are shown. (f) Rotarod performance on trial 1 for iSPN-Tsc1 WT, Het, and KO mice quantified as terminal speed. Bars represent mean +/− SEM. Dots represent individual mice (n is the same as in panel e). Kruskal-Wallis test, p=0.6800. (g) Overall learning rate of iSPN-Tsc1 WT, Het and KO mice calculated as the slope of the line of performance on the first trial to the last trial for individual mice (RPM/trial). Bars represent mean +/− SEM. Dots represent individual mice (n is the same as for panel e). Kruskal-Wallis test, p=0.2761. (h) Percentage of iSPN-Tsc1 WT, Het, and KO mice with a positive learning curve (slope of performance) from trial 7 to trial 12. Dark shaded regions indicate the percent of mice with a positive slope, light shaded regions indicate the percent of mice with a negative slope, within each genotype. The numbers on the bars indicate the percentage of mice of each genotype with a positive slope. N is the same as for panel e. (i) Performance on the accelerating rotarod across 12 trials (4 days) in germline *Tsc2^+/+^* (WT, n=17) and *Tsc2*^+/−^ (Het, n=31) mice. Circles represent mean +/− SEM. Mixed-effects analysis p values are shown. (j) Rotarod performance on trial 1 for Tsc2 WT and Het mice quantified as terminal speed. Bars represent mean +/− SEM. Dots represent individual mice (n is the same as in panel i). unpaired t-test, p=0.2533. (k) Overall learning rate of Tsc2 WT and Het mice calculated as the slope of the line of performance on the first trial (1) to the last trial (12) for individual mice (RPM/trial). Bars represent mean +/− SEM. Dots represent individual mice (n is the same as for panel i). Mann-Whitney test, **, p=0.0057. (l) Percentage of Tsc2 WT and Het mice with a positive learning curve (slope of performance) from trial 7 to trial 12. Dark shaded regions indicate the percent of mice with a positive slope, light shaded regions indicate the percent of mice with a negative slope, within each genotype. The numbers on the bars indicate the percentage of mice of each genotype with a positive slope. N is the same as for panel i.

To determine whether enhanced motor routine learning also occurs with brain-wide disruption of the Tsc1/2 complex, we tested mice that were heterozygous for a germline loss-of-function mutation in *Tsc2* (Onda et al., 1999). Heterozygous mice were used as germline homozygous deletion of *Tsc2* is embryonic lethal (Onda et al., 1999). Similar to dSPN-Tsc1 KO mice, *Tsc2* Het mice had normal initial motor performance on the accelerating rotarod but exhibited a significant increase in learning rate compared to WT littermates that became apparent in the more challenging trials of the test (Fig. 2i-l). Again, this was not likely due to a change in weight as this was not significantly different between genotypes (Tsc2 WT male = 27.41 +/− 0.63g vs Tsc2 Het male = 25.45 +/− 0.47g, p=0.0637, Mann-Whitney test; Tsc2 WT female = 19.53 +/− 0.55g vs Tsc2 Het female = 19.53 +/− 0.65g; p=0.9989, unpaired t-test). Together these results demonstrate that increased motor routine learning can be induced by even partial disruption of the Tsc1/2 complex and that loss of Tsc1 from dSPNs alone is sufficient to drive this change.

### Loss of Tsc1 increases cortical drive of direct pathway striatal neurons

Motor learning relies on corticostriatal transmission, therefore, changes in SPN response to cortical activity could account for enhanced motor routine learning in dSPN-Tsc1 KO mice. To test this, we crossed the *Tsc1^fl/fl^;Drd1-Cre;Ai9* and *Tsc1^fl/fl^;Adora2a-Cre;Ai9* mice to the *Thy1-ChR2-YFP* mouse line, which expresses channelrhodopsin (ChR2-YFP) in a subset of cortical layer V pyramidal cells (Arenkiel et al., 2007) (Fig. 3a). To simulate a train of cortical inputs, we applied ten light pulses over the recording site in dorsolateral striatum and recorded responses in SPNs in current-clamp mode in the absence of synaptic blockers (Fig. 3b). By varying the intensity of cortical stimulation, either subthreshold depolarizations or action potentials (APs) could be elicited in SPNs with a given probability. We quantified the number of APs evoked at different light intensities (Fig. S4) as a percentage of total stimulation events. We found that cortical terminal stimulation resulted in significantly increased spike probability across all light intensities in dSPN-Tsc1 KO cells compared to WT (Fig. 3c,d). Notably, loss of one copy of *Tsc1* in dSPNs was sufficient to increase corticostriatal drive as dSPN-Tsc1 Het neurons showed enhanced probability of cortically-driven spiking (Fig. 3c,d).

**Figure 3.**
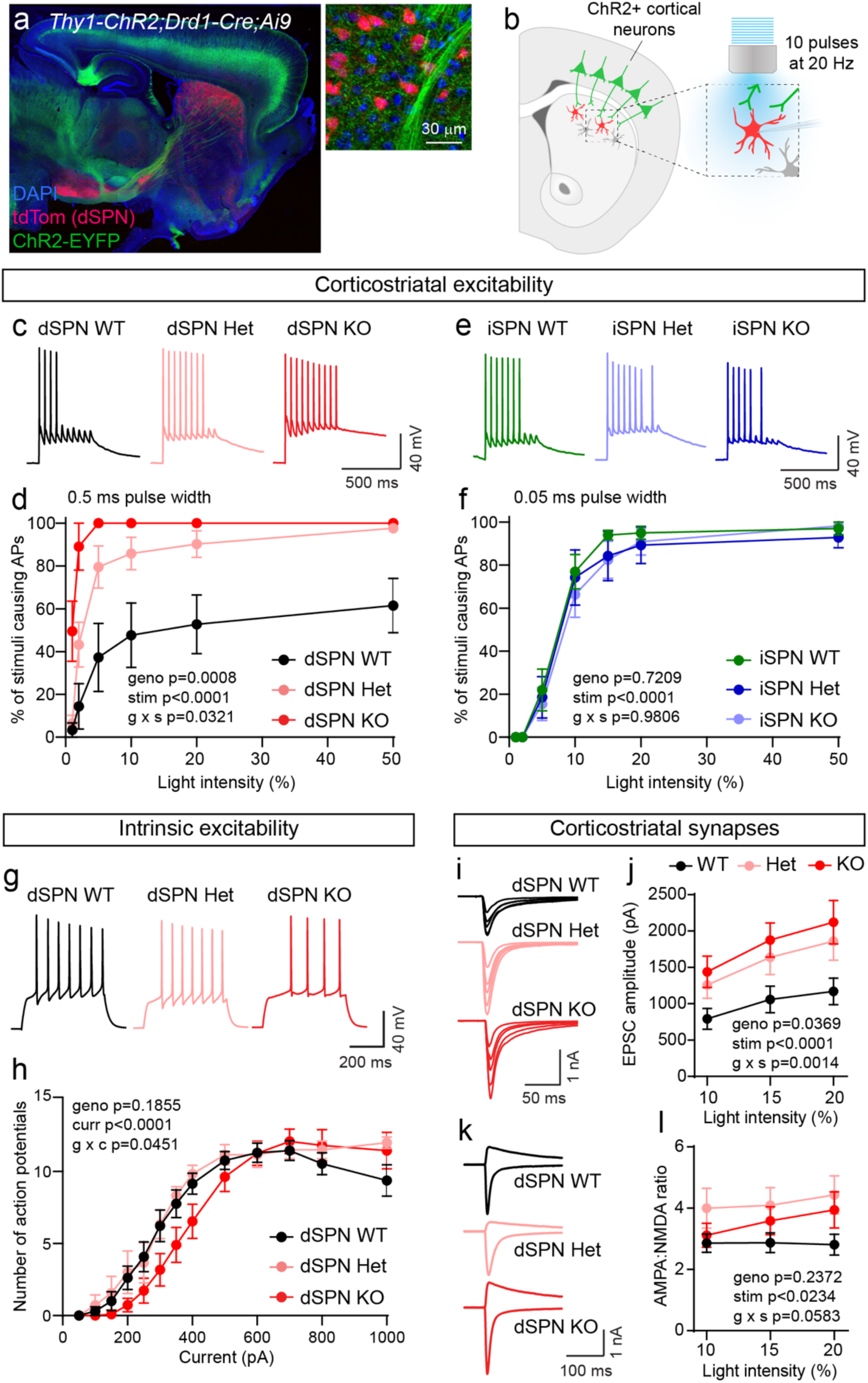
Tsc1 loss selectively increases cortico-dSPN synaptic drive. (a) Confocal image of a sagittal brain section from a *Tsc1^wt/wt^;Thy1-ChR2-EYFP*;*Drd1-Cre;Ai9* quadruple transgenic mouse. Right image shows cortical terminals expressing ChR2-EYFP in the dorsal striatum. (b) Schematic of the experiment. Cortical terminals expressing ChR2 were stimulated with 10 pulses of blue light at 20 Hz and responses were recorded in identified dSPNs in dorsolateral striatum. For dSPNs, the pulse width was 0.5 ms and for iSPNs the pulse width was 0.05 ms. (c,d) Example traces at 10% light intensity (c) and quantification as an input-output curve (mean +/− SEM) of action potentials (APs) evoked in dSPNs by cortical terminal stimulation (d) (*Tsc1;Thy1-ChR2-EYFP*;*Drd1-Cre;Ai9* mice). dSPN-Tsc1 WT n=5, dSPN Het n=7, dSPN KO n=5 neurons. Two-way ANOVA p values are shown. (e,f) Example traces at 10% light intensity (e) and quantification as an input-output curve (mean +/− SEM) of APs evoked in iSPNs by cortical terminal stimulation (f) (*Tsc1;Thy1-ChR2-EYFP*;*Adora2a-Cre;Ai9* mice). iSPN-Tsc1 WT n=10, dSPN Het n=11, dSPN KO n=7 neurons. Two-way ANOVA p values are shown. (g,h) Example traces from a 350 pA current step (g) and quantification (h) of the number of APs induced by depolarizing current steps in dSPN-Tsc1 WT, Het and KO neurons (*Tsc1;Drd1-Cre;Ai9* mice). Circles represent mean +/− SEM. dSPN WT n=13, dSPN Het n=6, dSPN KO n=11 neurons. Mixed-effects analysis p values are shown. (i) Example traces show EPSCs induced by corticostriatal stimulation at different light intensities (5-20%) in dSPN-Tsc1 WT, Het and KO neurons (*Tsc1;Thy1-ChR2-EYFP*;*Drd1-Cre;Ai9* mice). (j) Quantification of corticostriatal EPSC amplitude in dSPN WT, Het and KO neurons induced by different light intensities (0.5 ms pulse width). Circles represent mean +/− SEM. dSPN-Tsc1 WT n=18, dSPN Het n=7, dSPN KO n=25 neurons. Mixed-effects analysis p values are shown. (k) Example traces show pairs of EPSCs recorded at +40 mV (top traces) and −80 mV (bottom traces) from dSPN-Tsc1 WT, Het and KO neurons (*Tsc1;Thy1-ChR2-EYFP*;*Drd1-Cre;Ai9* mice). (l) Quantification of AMPA:NMDA ratio per cell in dSPN WT, Het and KO neurons across different intensities of corticostriatal stimulation. Circles represent mean +/− SEM. dSPN-Tsc1 WT n=9, dSPN Het n=7, dSPN KO n=14 neurons. Two-way ANOVA p values are shown.

We performed the cortical stimulation experiment in iSPNs and found that WT iSPNs were more readily driven to spike than dSPNs, consistent with their increased intrinsic excitability (Benthall et al., 2018; Gertler et al., 2008). Therefore, we reduced the length of the light pulse to avoid saturating the response. Interestingly, the number of APs elicited from iSPNs during cortical stimulation was not different between genotypes at any light intensity (Fig. 3e,f). These results are consistent with the rotarod results and demonstrate that deletion of *Tsc1* has a selective impact on dSPNs, while iSPNs are remarkably unaffected.

### Increased cortico-dSPN excitability results from enhanced synaptic transmission

Enhanced responses in dSPN-Tsc1 Het and KO mice to cortical terminal stimulation could result from increased intrinsic membrane excitability, increased synaptic excitation, or both. Indeed, *Tsc1* deletion has diverse effects on intrinsic and synaptic excitability depending on the cell type studied (Bateup et al., 2013; Benthall et al., 2018; Ehninger et al., 2008; Kosillo et al., 2019; Malik et al., 2019; Normand et al., 2013; Tavazoie et al., 2005; Tsai et al., 2012; Yang et al., 2012). To test if changes in intrinsic excitability occurred in dSPN-Tsc1 KO neurons, we injected positive current and measured the number of APs fired as a function of current step amplitude. The input-output curve for dSPN-Tsc1 KO cells was shifted slightly to the right relative to dSPN-Tsc1 WT and Het neurons, indicating a small decrease in intrinsic membrane excitability (Fig. 3g,h).

Intrinsic hypoexcitability has been observed in other neuron types with Tsc1 loss (Bateup et al., 2013; Kosillo et al., 2019; Normand et al., 2013; Tsai et al., 2012; Yang et al., 2012), and likely results from somatic hypertrophy (see Fig. 1), as membrane area and excitability are inversely related. However, this does not explain the enhanced corticostriatal excitability observed in dSPNs. To measure synaptic excitability, we recorded AMPA receptor (AMPAR)-mediated excitatory currents (EPSCs) evoked by optical stimulation of cortical terminals in *Tsc1^fl/fl^;Drd1-Cre;Ai9;Thy1-ChR2-YFP* mice. Stimulation of cortical terminals resulted in significantly larger AMPAR-mediated EPSCs in dSPN-Tsc1 Het and KO cells compared to WT cells, particularly at higher light intensities (Fig. 3i,j). When we measured the ratio of AMPAR currents recorded at −80 mV to NMDAR currents recorded at +40 mV, we found that there was a slight increase in the AMPA:NMDA ratio in dSPN-Tsc1 Het and KO neurons at higher light intensities; however, this was not statistically significant (Fig. 3k,l). These results suggest a global enhancement of excitatory synaptic function in Tsc1 KO dSPNs.

To investigate the specific synaptic properties that were altered in dSPN-Tsc1 Het and KO neurons, we recorded miniature excitatory postsynaptic currents (mEPSCs) at six weeks of age and found an increase in both amplitude and frequency of mEPSCs onto dSPN-Tsc1 KO cells compared to WT (Fig. 4a-c). Increased mEPSC frequency was also observed in dSPN-Tsc1 Het cells (Fig. 4a,c). To determine the developmental timing of these changes, we recorded mEPSCs in dSPN-Tsc1 WT and KO neurons at 2, 3 and 4 weeks of age. We found that the increased mEPSC amplitude and frequency in dSPN-Tsc1 KO neurons did not emerge until 4 weeks of age, a time when corticostriatal circuits are maturing and becoming refined (Kuo and Liu, 2019) (Fig. S5a,b). Thus, loss of Tsc1 may not affect the initial development of excitatory synapses but could affect their activity-dependent refinement. Consistent with a lack of cortico-iSPN excitability change, no significant changes in mEPSC amplitude or frequency were observed in iSPN-Tsc1 Het or KO cells at 6 weeks of age (Fig. 4d-f).

**Figure 4.**
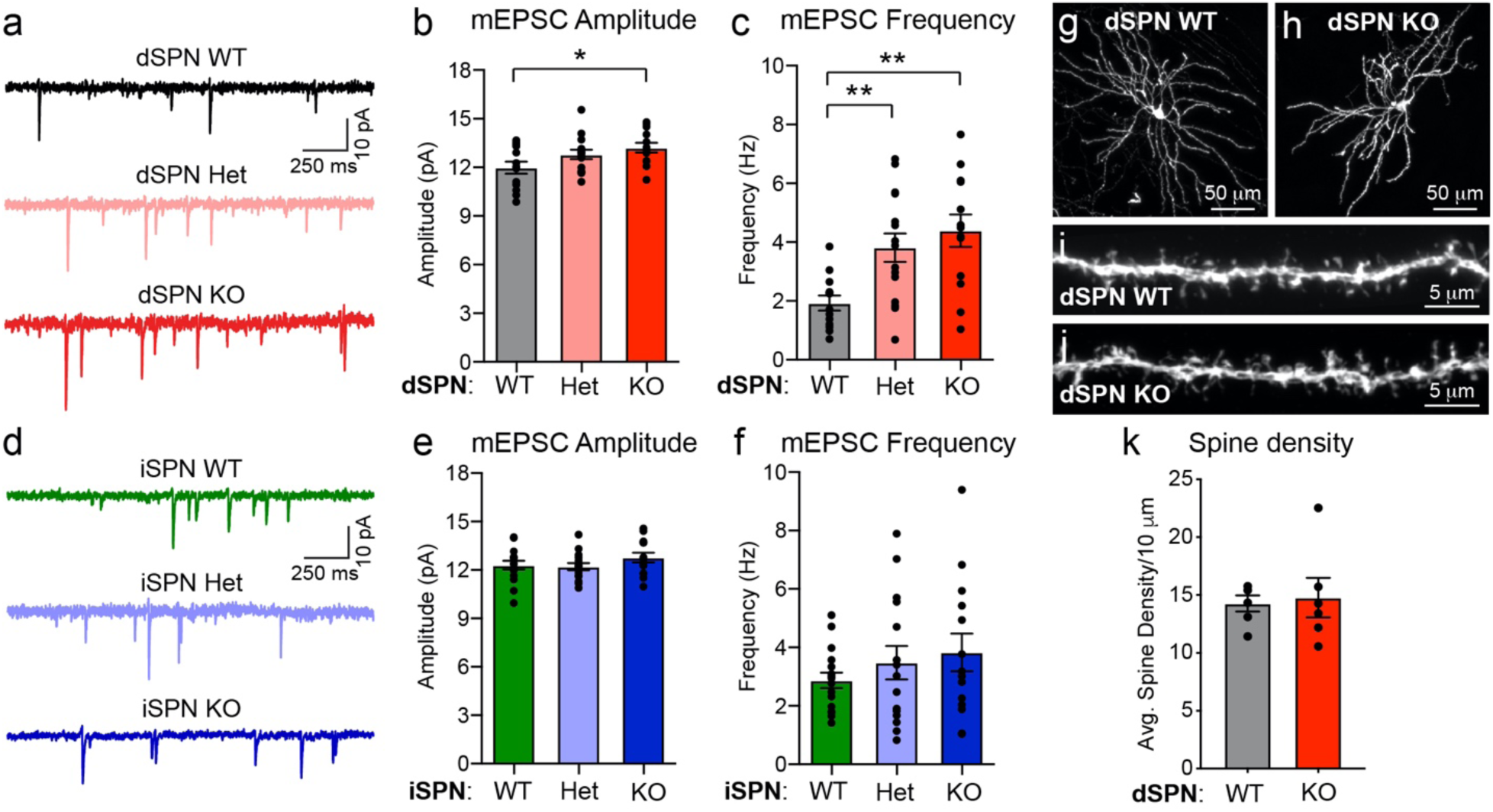
Loss of Tsc1 increases excitatory synaptic transmission onto dSPNs. (a) Example traces of miniature excitatory post-synaptic currents (mEPSCs) recorded from dSPN-Tsc1 WT, Het and KO neurons. (b,c) Mean +/− SEM mEPSC amplitude (b) and frequency (c) recorded from dSPN-Tsc1 WT, Het and KO neurons. Dots represent individual neurons. For panel b, one-way ANOVA, p=0.0337; Holm-Sidak’s multiple comparisons tests, dSPN WT vs Het, p=0.1346; dSPN WT vs KO, *, p=0.0204. For panel c, one-way ANOVA, p=0.0014; Holm-Sidak’s multiple comparisons tests, dSPN WT vs Het, **, p=0.0094; dSPN WT vs KO, **, p=0.0011. For both b and c, dSPN WT n=13, dSPN Het n=15, dSPN KO n=13 neurons. (d) Example traces of mEPSCs recorded from iSPN-Tsc1 WT, Het and KO neurons. (e,f) Mean +/− SEM mEPSC amplitude (e) and frequency (f) recorded from iSPN-Tsc1 WT, Het and KO neurons. Dots represent individual neurons. For panel e, one-way ANOVA, p=0.2942. For panel f, one-way ANOVA, p=0.4083. For both e and f, iSPN WT n=16, iSPN Het n=15, iSPN KO n=14 neurons. (g-h) Confocal images of individual Tsc1 WT (g) and KO (h) dSPNs labeled in *Tsc1;Drd1-Cre* mice using an AAV expressing Cre-dependent tdTomato. (i-j) Representative images of dendritic spines from a Tsc1 WT (i) and KO (j) dSPN. (k) Quantification of dendritic spine density in Tsc1 WT and KO dSPNs. Data are represented as mean +/− SEM spine density per 10 μm of dendrite. Dots display the average spine density for individual neurons. dSPN WT n=6 neurons (12 dendrites) from 3 mice, dSPN KO n=6 neurons (13 dendrites) from 3 mice. Mann-Whitney test, p=0.8182.

While changes in mEPSC amplitude usually reflect increased post-synaptic AMPAR function, increased mEPSC frequency can result from a greater number of synaptic contacts or a change in presynaptic release probability. To measure synapse number, we sparsely labeled dSPNs in the dorsal striatum using an AAV expressing Cre-dependent tdTomato. In fixed striatal sections, we imaged and reconstructed individual dSPNs and quantified the density of dendritic spines, which are the sites of corticostriatal synapses onto SPNs. We found that dSPN-Tsc1 KO neurons had equivalent spine density to dSPN-Tsc1 WT cells (Fig. 4g-k), suggesting that the increased mEPSC frequency was likely due to a change in presynaptic release probability.

### Endocannabinoid-mediated long-term depression is impaired in dSPN-Tsc1 KO neurons

One mechanism that could explain enhanced presynaptic corticostriatal transmission onto dSPN-Tsc1 KO neurons is a loss of long-term synaptic depression (LTD), which would render cells unable to depress excitatory inputs. Corticostriatal terminals express CB1 receptors that mediate endocannabinoid-LTD (eCB-LTD), a prominent form of striatal synaptic depression (Kreitzer and Malenka, 2008; Lovinger, 2010). Upon coincident activation of group 1 metabotropic glutamate receptors (mGluR1/5) and L-type calcium channels, SPNs release the endocannabinoids anandamide (AEA) or 2-arachidonoylglycerol (2-AG), which act as retrograde signals that activate presynaptic CB1Rs to decrease corticostriatal release probability (Luscher and Huber, 2010). It was previously thought that eCB-LTD occurs primarily in iSPNs (Kreitzer and Malenka, 2007), but recent studies using more selective stimulation of corticostriatal terminals have revealed eCB-LTD in both SPN subtypes (Wu et al., 2015). Notably, since another form of mGluR-dependent LTD expressed in the hippocampus is impaired in multiple TSC mouse models (Auerbach et al., 2011; Bateup et al., 2011; Chevere-Torres et al., 2012; Potter et al., 2013), we reasoned that loss of eCB-LTD could occur in dSPNs with *Tsc1* deletion.

We induced eCB-LTD in dSPNs with wash-on of the group 1 mGluR agonist DHPG and monitored the amplitude of EPSCs in response to optogenetic stimulation of cortical terminals in the dorsolateral striatum. While dSPN-Tsc1 WT neurons showed a long-lasting synaptic depression to ∼79% of baseline levels, eCB-LTD did not consistently occur in dSPN-Tsc1 KO neurons, which exhibited only a small initial reduction in EPSC amplitude during DHPG application that was not maintained (Fig. 5a). To ensure that the protocol was inducing eCB-LTD, we performed the DHPG wash-on in the presence of AM-251, a CB1R antagonist, and found that LTD was blocked in control dSPNs (Fig. S6). Loss of eCB-LTD in dSPN-Tsc1 KO neurons cannot be explained by a presynaptic difference in CB1R function, as direct activation of CB1Rs with the CB1R agonist WIN-2 depressed corticostriatal EPSCs to a similar extent in *Tsc1* deleted and control dSPNs (Fig. 5b).

**Figure 5.**
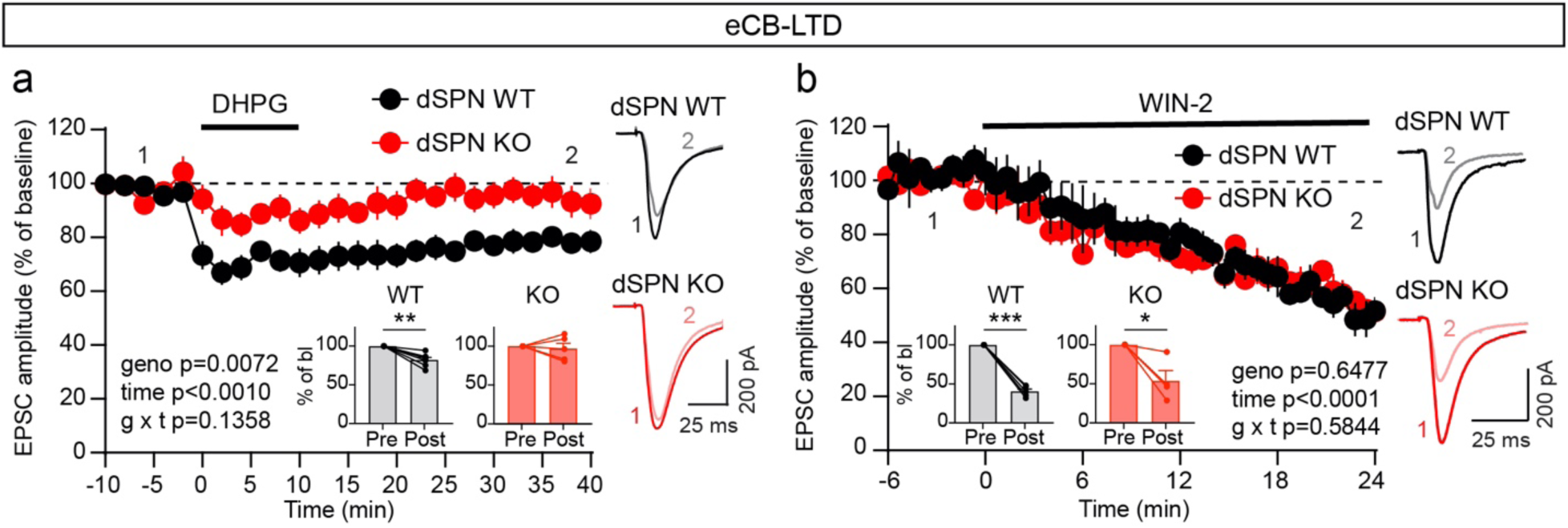
dSPN-Tsc1 KO neurons have impaired eCB-LTD. (a) Corticostriatal terminals in the dorsolateral striatum were stimulated with blue light (30 second ISI) to evoke EPSCs in dSPN-Tsc1 WT and KO neurons (*Tsc1;Thy1-ChR2-EYFP*;*Drd1-Cre;Ai9* mice). Following a 10-minute baseline period, DHPG (100 μM) was washed on for 10 minutes. Data are presented as mean +/− SEM percent of baseline. P values for mixed-effects analysis are shown. dSPN WT n=7, dSPN KO n=5 neurons. Inset graphs show the average (mean +/− SEM) EPSC during the 5-minute baseline preceding DHPG wash-on (Pre) and 35-40 minutes after DHPG wash-on (Post) expressed as a percentage of the baseline response. Dots represent individual cells. dSPN WT, **, p=0.0014; dSPN KO, p=0.6896, paired t-tests. Example traces on the right show average EPSCs from the baseline period (“1”, dark lines) and 35-40 minutes after DHPG application (“2”, light lines). (b) Corticostriatal terminals were stimulated with blue light (30 second ISI) to evoke EPSCs in dSPN-Tsc1 WT and KO neurons (*Tsc1;Thy1-ChR2-EYFP*;*Drd1-Cre;Ai9* mice). WIN-2 (2 μM) was applied at time 0 to activate CB1 receptors. Data are presented as mean +/− SEM percent of baseline. P values for mixed-effects analysis are shown. dSPN WT n=5, dSPN KO n=4 neurons. Insets show the average (mean +/− SEM) EPSC during the 3-minute baseline preceding WIN-2 wash-on (Pre) and 21-24 minutes after WIN-2 wash-on (Post) expressed as a percentage of the baseline response. Dots represent individual cells. dSPN WT, ***, p<0.0001; dSPN KO, *, p=0.0389, paired t-tests. Example traces on the right show the average EPSC from the last 5 minutes of the baseline period (“1”, dark lines) and 19-24 minutes after WIN-2 application (“2”, light lines).

To investigate a potential molecular basis for the loss of eCB-LTD in Tsc1 KO dSPNs, we performed translating ribosome affinity purification (TRAP) to assess the translational status of mRNAs encoding key proteins in the eCB-LTD pathway in dSPNs. To do this, we engineered an AAV to express a Cre-dependent GFP-tagged ribosomal protein (EGFP-L10a) and injected this into the dorsal striatum of *Drd1-Cre;Ai9* mice that were WT or homozygous floxed for *Tsc1* (Fig. 6a,b). This approach enabled selective expression of EGFP-L10a in dSPNs (Fig. 6c). Using EGFP as an affinity tag, we isolated ribosome-bound mRNAs selectively from dSPNs using TRAP (Heiman et al., 2014; Heiman et al., 2008). We verified the specificity of this approach using qPCR to quantify the relative levels of *Drd1* (D1 receptor) and *Drd2* (D2 receptor) mRNA in the TRAP samples compared to the unbound samples, which includes RNA from all striatal cell types. As expected, *Drd1* mRNA was significantly enriched in the TRAP samples compared to the unbound samples (Fig. 6d). Accordingly, *Drd2* mRNA was depleted from the TRAP samples, demonstrating preferential isolation of mRNA from dSPNs (Fig. 6e).

**Figure 6.**
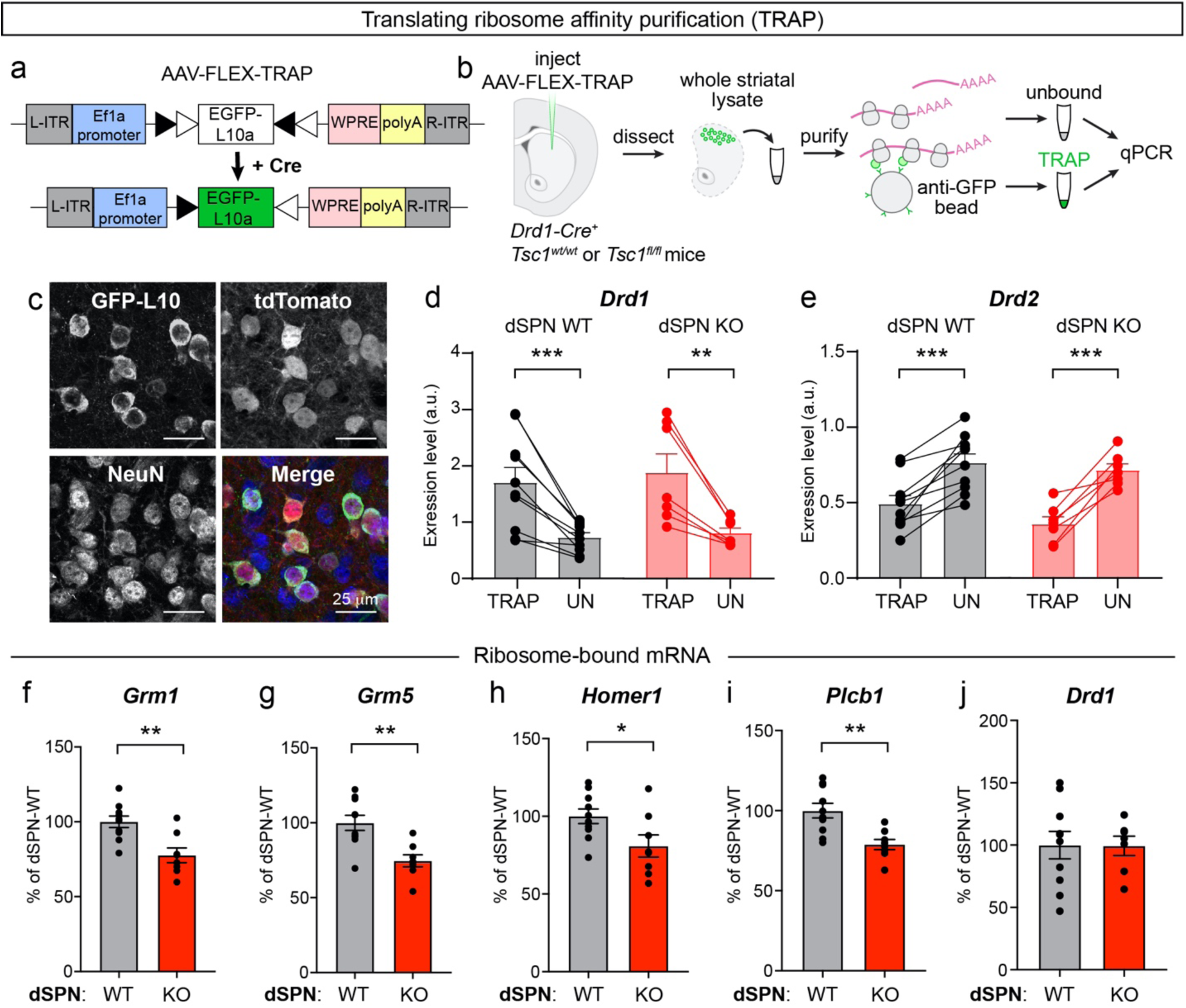
dSPN Tsc1 KO cells exhibit reduced ribosome engagement of mRNAs involved in eCB-LTD. (a) Schematic of the design of AAV-FLEX-TRAP, based on a double-floxed inverted open reading frame (DIO) which allows Cre-dependent expression of a GFP-tagged ribosomal subunit (EGFP-L10a). (b) Schematic of the workflow for the Translational Ribosome Affinity Purification (TRAP) experiment. (c) Confocal image of EGFP-L10a expression from an AAV5-hSyn-DIO-EGFP-L10a-WPRE-hGH injection into the dorsolateral striatum of a *Tsc1^fl/fl^;Drd1-Cre;Ai9* mouse showing expression in tdTomato+ dSPNs. (d-e) Quantification of relative *Drd1* (d) and *Drd2* (e) mRNA levels in TRAP samples (ribosome bound mRNA isolated from dSPNs) versus the unbound (all striatal tissue) from the same mouse measured by qPCR. Bars represent mean +/− SEM. Dots represent individual samples, taken from one mouse. For panel d, ***, p=0.0009 (dSPN WT); **, p=0.0053 (dSPN KO), paired t-tests. For panel e, ***, p=0.0003 (dSPN WT); ***, p=0.0007 (dSPN KO), paired t-tests. n=10 dSPN WT and 7 dSPN KO mice. (f-j) Quantification of *Grm1* (f), *Grm5* (g), *Homer1* (h), *Plcb1* (i), and *Drd1* (j) mRNA levels in TRAP samples (ribosome bound mRNA isolated from dSPNs) relative to *Actb* from dSPN-Tsc1 WT and KO mice measured by qPCR. Bars represent mean +/− SEM and dots represent individual mice. For panel f, **, p=0.0022; panel g, **, p=0.0018; panel h, *, p=0.0346; panel i, **, p=0.0025; panel j, p=0.9702; unpaired t-tests. For all panels, dSPN WT n=10 and dSPN KO n=8 mice (except for *Drd1* dSPN KO, n=7 mice).

We compared the relative expression levels of mRNAs encoding proteins required for eCB-LTD including the two types of group 1 mGluRs (*Grm1* and *Grm5*), the requisite mGluR scaffold protein Homer 1 (*Homer1*), and phospholipase C beta (*Plcb1*), which promotes the formation of diacylglycerol that is subsequently converted to 2-AG by diacylglycerol lipase (Ohno-Shosaku and Kano, 2014). We also compared the levels of ribosome-bound *Drd1* as a non-eCB-LTD control. We found that dSPN-Tsc1 KO mice had increased amounts of total ribosome-bound mRNA (0.0518 +/− 0.0077 vs 0.0313 +/− 0.0031 ratio of TRAP-isolated mRNA to unbound RNA in dSPN KO vs WT mice, p=0.0138, unpaired t-test), consistent with a global increase in protein synthesis resulting from high mTORC1 activity. However, all four components of the eCB-LTD pathway showed a relative reduction in ribosome-bound mRNA in dSPN-Tsc1 KO mice compared to WT mice when normalized to *Actb* (B-Actin) (Fig. 6f-i). Notably, levels of *Drd1* were not significantly different between genotypes (Fig. 6j). The amount of ribosome-bound mRNA is generally thought to reflect translation efficiency (Gobet and Naef, 2017), therefore these results suggest a relative downregulation in the translation of multiple mRNAs encoding proteins required for eCB-LTD in Tsc1 KO dSPNs. Such decreased expression may be one mechanism responsible for the lack of functional eCB-LTD in Tsc1 KO dSPNs.

## Discussion

In this study we tested whether cell type-specific deletion of *Tsc1* from striatal neurons was sufficient to alter synaptic function and motor behaviors. We find that in both direct and indirect pathway SPNs, developmental loss of *Tsc1* upregulates mTORC1 signaling and increases soma size, consistent with other neuron types. However, we find that mTORC1 activation only in dSPNs, but not iSPNs, enhances motor routine learning in the absence of spontaneous stereotypies or locomotor hyperactivity. Further, we find that loss of Tsc1 from dSPNs is associated with increased corticostriatal synaptic excitability and an impairment in eCB-LTD. Notably, loss of only one copy of *Tsc1* is sufficient to increase cortical drive of dSPNs and enhance motor learning. These findings implicate the striatal direct pathway as a possible driver of altered motor behaviors in TSC and identify a potential synaptic mechanism for this.

It has been shown that loss of Tsc1 in either the cerebellum or thalamus is sufficient to cause social behavior deficits and spontaneous RRBs including repetitive self-grooming in TSC mouse models (Normand et al., 2013; Tsai et al., 2012). Given the central role of the striatum in action selection and motor learning, we hypothesized that alterations in striatal circuits might also contribute to altered motor behaviors in the context of TSC. We found that dSPN or iSPN-specific *Tsc1* deletion was not sufficient to induce spontaneous repetitive behaviors, suggesting that other brain regions may be more important for initiating these behaviors. Indeed, the striatum has been shown to be important for the proper sequencing of grooming action patterns but not the initiation of grooming bouts (Kalueff et al., 2016). In line with this, it was reported that striatal-specific loss of the ASD-risk gene *Shank3* was not sufficient to reproduce the repetitive self-grooming that is observed with global loss of *Shank3* but that this could be driven by loss of *Shank3* from forebrain excitatory neurons (Bey et al., 2018). Alternatively, the central striatum was recently shown to be the key area for the initiation of self-grooming, and in our model, *Tsc1* was primarily deleted from dSPNs in the dorsal striatum (Corbit et al., 2020).

Intriguingly, we found that mice with loss of Tsc1 in dorsal dSPNs had enhanced performance on the accelerating rotarod, a motor learning task that relies on corticostriatal transmission and SPN plasticity (Yin et al., 2009). Previous work has shown that striatal mTORC1 signaling is required for motor learning on the accelerating rotarod, as both pharmacological and genetic intrastriatal inhibition of mTOR signaling impairs rotarod learning (Bergeron et al., 2014). Our results are congruent with these findings and show that the mTORC1-dependence of motor learning is bidirectional, as increasing mTORC1 activity in dSPNs enhances performance. Notably, an extended accelerating rotarod task may be necessary to reveal motor learning phenotypes, as *Tsc1* and *Tsc2* heterozygous mice were reported to have no changes in rotarod performance when an abbreviated, less challenging version of the task was used (Pryor et al., 2014; Sato et al., 2012). Importantly, the accelerating rotarod task has recently been recognized as commonly affected in mice with mutations in ASD-risk genes, as at least seven other genetic ASD mouse models have enhanced rotarod performance (Chadman et al., 2008; Hisaoka et al., 2018; Kwon et al., 2006; Nakatani et al., 2009; Penagarikano et al., 2011; Platt et al., 2017; Rothwell et al., 2014) (although this is not the case for all ASD-risk genes, see (Portmann et al., 2014; Sonzogni et al., 2018; Wang et al., 2016)). Most of the aforementioned mouse models have global gene deletions in which brain regions and circuits outside of the striatum could contribute to the phenotype. However, two studies showed that selectively disrupting ASD-risk genes (*Nlgn3* or *Chd8*) in striatal neurons was sufficient to increase rotarod learning (Platt et al., 2017; Rothwell et al., 2014). Moreover, similar to what was observed here, loss of *Nlgn3* from dSPNs alone led to enhanced motor learning, implicating the direct pathway as a key driver of this phenotype (Rothwell et al., 2014). Interestingly, for *Nlgn3* and *Chd8*, ventral striatal disruption of the gene was responsible for the motor learning improvement. Here, deletion of *Tsc1* from dSPNs was largely restricted to the dorsal striatum, and our findings are consistent with literature establishing the dorsal striatum’s role in motor learning (Yin et al., 2009). Further studies will be needed to define the contributions of dorsal versus ventral striatal circuits to motor learning and elucidate the contributions of specific striatal subregions to behavioral changes in TSC mouse models.

While enhanced motor routine learning may be a shared phenotype across multiple mouse models of ASD, the synaptic and cellular mechanisms driving this phenotype are likely to be distinct depending on the specific gene that is altered. dSPN-Nlgn3 KO mice displayed normal excitatory synaptic transmission and eCB-LTD but had a deficit in synaptic inhibition (Rothwell et al., 2014). In the global *Chd8^+/−^* mouse model, there was increased spontaneous excitatory transmission onto SPNs; however, the SPN sub-type was not defined (Platt et al., 2017). Here we found that dSPN-Tsc1 Het and KO cells had strongly enhanced corticostriatal excitation that was due to increased synaptic, but not intrinsic, excitability. We have previously reported increased presynaptic release probability in dSPNs with acute loss of Tsc1 (Benthall et al., 2018) and consistent with this, we found increased mEPSC frequency in the absence of changes to spine density in dSPN-Tsc1 KO neurons. Given that the cortical inputs were wild-type in our model, a potential explanation for increased mEPSC frequency is a change in retrograde signaling from dSPNs to cortical terminals. Indeed, we found that eCB-LTD was absent in Tsc1 KO dSPNs despite normal presynaptic CB1 receptor function. This suggests that loss of Tsc1 and upregulation of mTORC1 signaling in dSPNs interferes with one or more post-synaptic processes required for this form of plasticity: 1) expression of group 1 mGluRs, 2) signaling downstream of mGluRs, and/or 3) synthesis and release of eCBs. In support of this idea, we found that multiple mRNAs encoding proteins required for eCB-LTD exhibited differential ribosome engagement in dSPN-Tsc1 KO cells compared to WT and were relatively reduced. Together, this suggests that loss of Tsc1 and deregulation of mTORC1 signaling in dSPNs impairs post-synaptic mGluR signaling via altered expression of key components of this pathway.

A chronic impairment of eCB-LTD in Tsc1 KO dSPNs could explain not only the increased mEPSC frequency but potentially the increased amplitude, as these synapses may have a greater capacity to undergo long-term potentiation, leading to synaptic strengthening over time. This idea is consistent with our observation that changes in mEPSC properties in dSPN-Tsc1 KO neurons do not arise until four weeks of age and later, after the primary period of postnatal striatal maturation (Lieberman et al., 2018). A causal role for impaired eCB-LTD in increased AMPAR function is also suggested by the finding that direct disruption of endocannabinoid synthesis and release from dSPNs is sufficient to increase their glutamatergic drive (Shonesy et al., 2018). Interestingly, this study showed that disruption of 2-AG signaling in dSPNs had behavioral consequences while loss from iSPNs had no measurable effects on behavior (Shonesy et al., 2018), consistent with the lack of behavior changes reported here for iSPN-Tsc1 KO mice.

Taken together, our results support a model whereby developmental loss of Tsc1 from dSPNs impairs eCB-LTD resulting in unchecked corticostriatal drive. While further work will be needed to establish a causal link between enhanced cortico-dSPN activity and increased rotarod learning, it is possible that perturbed presynaptic plasticity in Tsc1 KO dSPNs may alter corticostriatal coupling during rotarod training, leading to atypical learning in this paradigm. In terms of autistic behaviors in individuals with TSC, our findings suggest that striatal synaptic and circuit changes, which in mouse models increase their ability to learn a stereotyped motor routine, could be a contributor to the emergence of restricted, repetitive patterns of behavior.

## Acknowledgements

This work was supported by a SFARI Pilot Award (# 307866), R21 NS096415, R56 MH111821 and R01 NS105634 grants to H.S.B. H.S.B. is a Chan Zuckerberg Biohub Investigator and was supported by an Alfred P. Sloan Foundation fellowship. K.N.B. was supported by an NSF graduate research fellowship (DGE 1106400). We thank Corinna Wong and Caroline Keeshen for their assistance with mouse colony maintenance. We thank Dr. John Blair for assistance with cloning the AAV-FLEX-TRAP construct. We thank Dr. Claire Darzacq for access to the qPCR equipment. We thank Dr. Anne Schaefer for advice on the TRAP protocol and for providing the EGFP-L10a construct. We thank Kamran Ahmed for generating the figure schematics. Confocal spine imaging experiments were conducted at the CRL Molecular Imaging Center, supported by the Helen Wills Neuroscience Institute. We thank Holly Aaron and Feather Ives for their microscopy training and assistance.

## Author Contributions

Conceptualization, H.S.B.; Methodology, K.N.B.; Formal Analysis, K.N.B., K.R.C., A.H.C.W.A-M., E.Y.C., and H.S.B.; Investigation, K.N.B., K.R.C., A.H.C.W.A-M., and E.Y.C.; Writing-Original Draft, K.N.B. and H.S.B.; Writing-Review & Editing, K.N.B., K.R.C., and H.S.B.; Visualization, K.N.B. and H.S.B.; Supervision, K.N.B. and H.S.B.; Funding Acquisition, H.S.B.

## Methods

### Mouse lines

Animal experiments were performed in accordance with protocols approved by UC Berkeley’s Animal Care and Use Committee. Male and female mice were used for all experiments. The ages used are indicated below for each experiment. All mice used for experiments were heterozygous or hemizygous for the *Ai9*, *Drd1-Cre*, *Adora2a-Cre*, or *Thy1-ChR2-YFP* transgenes to avoid potential physiological or behavioral alterations.

The following mouse lines were used:

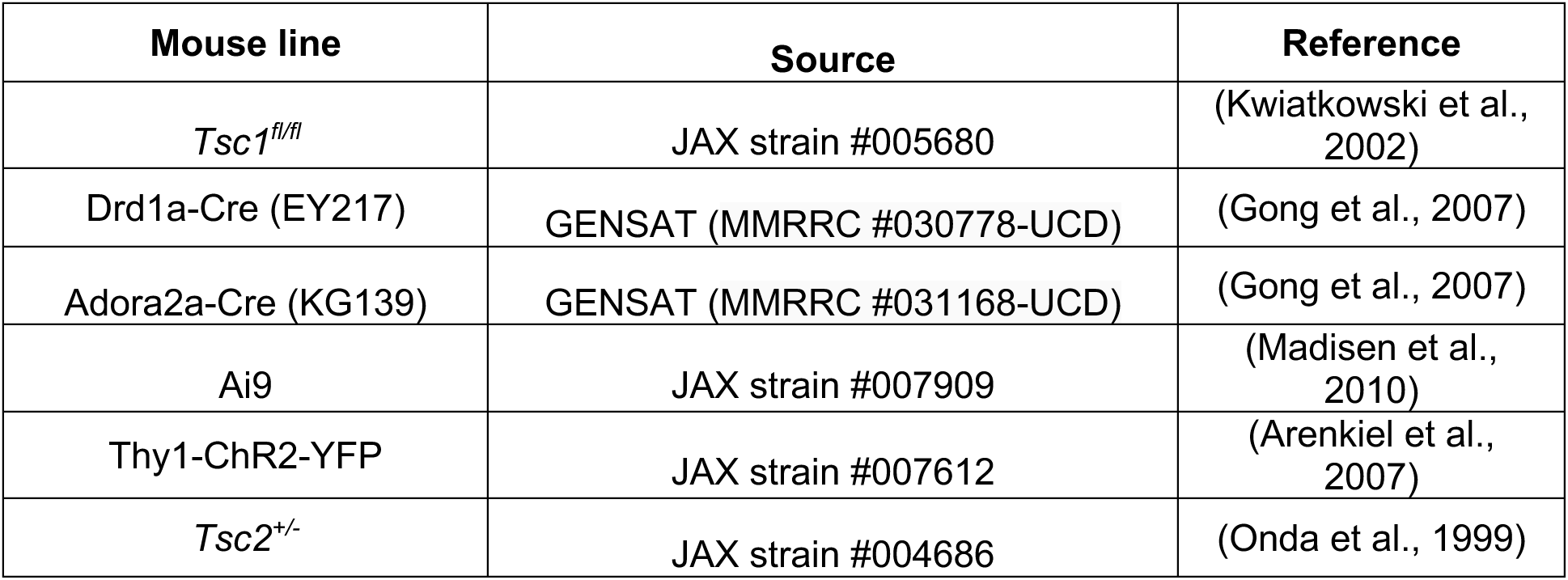

| Mouse line | Source | Reference |
| --- | --- | --- |
| *Tsc1<sup>fl/fl</sup>* | JAX strain #005680 | (Kwiatkowski et al., 2002) |
| Drd1a-Cre (EY217) | GENSAT (MMRRC #030778-UCD) | (Gong et al., 2007) |
| Adora2a-Cre (KG139) | GENSAT (MMRRC #031168-UCD) | (Gong et al., 2007) |
| *Ai9* | JAX strain #007909 | (Madisen et al., 2010) |
| Thy1-ChR2-YFP | JAX strain #007612 | (Arenkiel et al., 2007) |
| *Tsc2<sup>+/-</sup>* | JAX strain #004686 | (Onda et al., 1999) |

### Immunohistochemistry

Mice (P40-50) were perfused transcardially with 1x PBS and 4% paraformaldehyde. Brains were post-fixed in 4% paraformaldehyde overnight, cryoprotected in sucrose, and sectioned coronally at 30 μm. Sections were blocked for 1 hr at room temperature (RT) in BlockAid (Thermo Fisher Scientific) and incubated for 48 hr at 4°C with antibodies against phosphorylated S6 ribosomal protein (Ser240/244, 1:800, Cell Signaling Technology, catalog # 5364S), GFP (1:1000, Abcam, catalog # ab13970) and NeuN (1:800, Millipore, catalog # MAB377). Sections were washed and incubated for 1 h at RT with Alexa Fluor 633- and 488-conjugated secondary antibodies (1:500, Invitrogen, catalog #’s A-21070 and A-31553). Sections were mounted onto slides using Vectashield reagent with DAPI (Vector Labs).

### Confocal imaging and analysis

To analyze p-S6 levels and soma volume, Z-stack images of dorsal striatal sections were taken on a confocal microscope (Zeiss LSM 780 AxioExaminer or Olympus FLUOVIEW FV3000) with a 20x objective using the same acquisition settings for each section. For quantification, cellular regions of interest (ROIs) were automatically created based on the tdTomato signal with the Surfaces module in Imaris software (Oxford Instruments). The mean p-S6 fluorescence intensity per ROI and average soma volume were calculated using Imaris. Values for Tsc1 Het and KO cells were normalized to the average of all wild-type cells imaged in the same batch. To generate cumulative probability plots, 300 cells from each mouse were used (100 or 150 cells per slice, 2-3 slices per mouse).

To analyze Cre-mediated recombination patterns in *Drd1*- and *Adora2a-Cre;Ai9* mice, Z-stack images of dorsolateral striatum, dorsomedial striatum, nucleus accumbens lateral shell, and nucleus accumbens medial shell were taken on a confocal microscope (Olympus FLUOVIEW FV3000) with a 20x objective. For quantification, ROIs were manually defined in ImageJ for all NeuN positive cells and used to determine percent co-localization with the Cre-dependent tdTomato reporter.

To analyze AAV-FLEX-TRAP (EGFP-L10a) expression, Z-stack images of dorsal striatum were taken on a confocal microscope (Olympus FLUOVIEW FV3000) with a 20x objective. For quantification, ROIs were manually defined in ImageJ for all GFP positive cells and used to verify co-localization with the Cre-dependent tdTomato reporter. 85-93% of EGFP-L10a positive cells in the dorsal striatum were tdTomato positive.

### Dendritic spine imaging and analysis

Neonatal (P1-4) dSPN-Tsc1 KO mice were cryoanesthetized and injected bilaterally with 200 nL AAV9.CAG.Flex.tdTomato.WPRE.bGH (AllenInstitute864), diluted 1:500 in sterile saline to achieve sparse transduction. Injections were targeted to the striatum, with coordinates approximately 1.2 mm lateral to midline, 2.0 mm posterior to bregma, and 1.5 mm ventral to the head surface. At age P40-50, mice were perfused and brains were post-fixed with 4% paraformaldehyde, then sectioned at 80 μm. Sections were blocked for 1 hr at RT in BlockAid (Thermo Fisher Scientific) and incubated for 48 hr at 4°C with an antibody against RFP (1:1000, Rockland (VWR), catalog #RL600-401-379). Sections were washed and incubated for 1 hr at RT with Alexa Fluor 546 secondary antibody (1:500, Invitrogen, catalog #A-11035). Sections were mounted onto slides using Vectashield reagent with DAPI (Vector Labs). Z-stack images of individual dendrites were taken on a confocal microscope (Zeiss LSM 880 NLO AxioExaminer) with a 63x objective using Airyscan super-resolution imaging. To quantify spine density, dendrites and spines were reconstructed using the FilamentTracer module in Imaris software (Oxford Instruments). The spine density of each dendrite was calculated using Imaris. Spine density analysis was initially separated into the first 40 μm of the dendrite (proximal) and the final 40 μm of the dendrite (distal). There was no significant difference between proximal and distal spine density within cells or across genotypes. These values were therefore combined and spine density of the entire 80 μm length of dendrite was reported.

### Behavioral analysis

Behavior studies were carried out in the dark phase of the light cycle under red lights (open field) or white lights (rotarod). Mice were habituated to the behavior testing room for at least 30 min prior to testing and covered by a black-out curtain. Mice were given at least one day between different tests. All behavior equipment was cleaned between each trial and mouse with 70% ethanol, and additionally rinsed in diluted soap followed by water at the end of the day. If male and female mice were to be tested on the same day, male mice were run first then returned to the husbandry room, after which all equipment was thoroughly cleaned prior to bringing in female mice for habituation. All animals to be tested from a given cage were run in each behavior test in the same day. Behavioral tests were performed with young adult male and female mice (6-10 weeks old). Mice had access to a running wheel in their home cage. The experimenter was blind to genotype throughout the testing and scoring procedures.

#### Open field

Exploratory behavior in a novel environment and general locomotor activity were assessed by a 60 min session in an open field chamber (40 cm L x 40 cm W x 34 cm H) made of transparent plexiglass. Horizontal infrared photobeams were positioned to detect rearing. The mouse was placed in the bottom right hand corner of the arena and behavior was recorded using an overhead camera and analyzed using the ANY-maze (Stoelting Co.) behavior tracking software. An observer manually scored self-grooming behavior during the first 20 minutes of the test. A grooming bout was defined as an unbroken series of grooming movements, including licking of body, paws, or tail, as well as licking of forepaws followed by rubbing of face with paws.

#### Rotarod

The accelerating rotarod test was used to examine motor learning. Mice were trained on a rotarod apparatus (Ugo Basile: 47650) for four consecutive days. Three trials were completed per day with at least a 5 min break between trials. The rotarod was accelerated from 5-40 revolutions per minute (rpm) over 300 s for trials 1-6 (days 1 and 2), and from 10-80 rpm over 300 s for trials 7-12 (days 3 and 4). On the first testing day, mice were acclimated to the apparatus by being placed on the rotarod rotating at a constant 5 rpm for 60 s and returned to their home cage for 5 minutes prior to starting trial 1. Latency to fall, or to rotate off the top of the rotarod barrel, was measured by the rotarod stop-trigger timer.

### Slice preparation for electrophysiology

Mice (P40-50) were perfused transcardially with ice-cold ACSF (pH=7.4) containing (in mM): 127 NaCl, 25 NaHCO_3_, 1.25 NaH_2_PO_4_, 2.5 KCl, 1MgCl_2_, 2 CaCl_2_, and 25 glucose, bubbled continuously with carbogen (95% O_2_ and 5% CO_2_). Brains were rapidly removed and coronal slices (275 μm) were cut on a VT1000S vibrotome (Leica) in oxygenated ice-cold choline-based external solution (pH=7.8) containing (in mM): 110 choline chloride, 25 NaHCO_3_, 1.25 NaHPO_4_, 2.5 KCl, 7 MgCl_2_, 0.5 CaCl_2_, 25 glucose, 11.6 sodium ascorbate, and 3.1 sodium pyruvate. Slices were recovered in ACSF at 34°C for 15 min and then kept at RT before recording.

### Whole-cell patch-clamp recording

Recordings were made with a MultiClamp 700B amplifier (Molecular Devices) at RT using 3-5 MOhm glass patch electrodes (Sutter, BF150-86-7.5). Data was acquired using ScanImage software, written and maintained by Dr. Bernardo Sabatini (https://github.com/bernardosabatini/SabalabAcq). Traces were analyzed in Igor Pro (Wavemetrics). Recordings with a series resistance >25 MOhms or holding current above −200 pA were rejected.

#### Current-clamp recordings

Current clamp recordings were made using a potassium-based internal solution (pH=7.4) containing (in mM): 135 KMeSO_4_, 5 KCl, 5 HEPES, 4 Mg-ATP, 0.3 Na-GTP, 10 phosphocreatine, and 1 ETGA. For corticostriatal excitability experiments, optogenetic stimulation consisted of a full-field pulse of blue light (470 nm, 0.15 ms for iSPNs or 0.5 ms for dSPNs, CoolLED) through a 63x objective. Light power was linear over the range of intensities tested (Fig. S3). No synaptic blockers were included. For intrinsic excitability experiments, NBQX (10 μM, Fisher, cat. #104410), CPP (10 μM, Fisher, cat. #24710) and picrotoxin (50 μM, Abcam, cat. #120315) were added to the external solution to block synaptic transmission. 500 ms depolarizing current steps were applied to induce action potentials. No holding current was applied to the membrane.

#### Voltage-clamp recordings

Voltage-clamp recordings were made using a cesium-based internal solution (pH=7.4) containing (in mM): 120 CsMeSO_4_, 15 CsCl, 10 TEA-Cl, 8 NaCl, 10 HEPES, 1 EGTA, 5 QX-314, 4 Mg-ATP, and 0.3 Na-GTP. Recordings were acquired with the amplifier Bessel filter set at 3 kHz. Miniature excitatory synaptic currents (mEPSCs) were recorded in the presence of TTX (1 μM, Abcam, cat. #120055) to prevent action potential-mediated release. Picrotoxin (50 μM, Abcam, cat. #120315) and CPP (10 μM, Fisher, cat. #24710) were included for mEPSC experiments to isolate AMPAR-mediated events. Corticostriatal synaptic stimulation experiments to measure evoked AMPA-mediated EPSCs were performed in picrotoxin (50 μM) and CPP (10 μM), and optogenetic stimulation consisted of a full-field pulse of blue light (470 nm, 0.15 ms) through a 63x objective. To measure AMPA/NMDA ratio, experiments were performed in 50 μM picrotoxin and the membrane was held at different potentials to isolate primarily AMPAR (−80 mV) or compound AMPAR and NMDAR (+40 mV) currents. The current at +40 mV was measured 50 ms after stimulation, by which time the AMPAR-mediated current has decayed.

#### eCB-LTD

Endocannabinoid-mediated long-term depression (eCB-LTD) was induced in *Thy1-ChR2;Drd1-Cre;Ai9;Tsc1* mice by bath application of the group 1 mGluR agonist DHPG (100 μM, Sigma, cat. #D3689) for 10 min, following a 10 min baseline measurement of EPSC amplitude with single full field light pulses (3-15% light intensity, 0.15 ms) delivered every 30 seconds to stimulate corticostriatal terminals. Light intensity was adjusted for each cell to evoke 500-700pA currents during the baseline period. DHPG was subsequently washed off and EPSC amplitude was monitored every 30 sec for an additional 40 min. Picrotoxin (50 μM) was added to the bath during eCB-LTD experiments to isolate excitatory events, and perfusion flow rate was set to 5 mL/min. Cells were held at −50 mV to facilitate opening of L-type calcium channels. CB1R antagonist AM-251 (10 μM, Fisher, cat. #111710) was added to the bath during a subset of eCB-LTD experiments with dSPN-Tsc1 WT cells to verify that the LTD observed during these experiments was dependent upon CB1R activation. For CB1R agonism experiments, WIN2 (2 μM, EMD Millipore, cat. #5.04344.0001) was applied to the bath throughout the recording, following a baseline period.

### AAV-Flex-TRAP plasmid and virus construction

The AAV-hSyn-DIO-EGFP-L10a-WPRE-hGH and AAV-Ef1a-DIO-EGFP-L10a-WPRE-hGH plasmids were assembled from pAAV-hSyn-DIO-mCherry (Addgene plasmid #50459) and pAAV-EF1A-DIO-mCherry (Addgene plasmid #50462), which were gifts from Bryan Roth, and an EGFP-L10a construct, which was a gift from Anne Schaefer. AAV5 viruses were prepared by the University of Pennsylvania Vector Core with a titer of 5.97 x 10^12^ for AAV5-hSyn-DIO-EGFP-L10a-WPRE-hGH and 3.22 x 10^12^ for AAV5-Ef1a-DIO-EGFP-L10a-WPRE-hGH.

### Intracranial virus injection

Mice were anesthetized with isoflurane and mounted on a stereotaxic frame equipped with ear cups. 800 nl of an AAV serotype 5 *Ef1a* or *hSyn* promoter-driven DIO-EGFP-L10a virus (AAV-FLEX-TRAP) was bilaterally injected into the dorsal striatum of 3-8 week old *Tsc1^cKO^;Drd1a-Cre(EY217)* mice. Coordinates for injection were +/−1.6 M/L, +0.6 A/P, −1.3 D/V. Mice were used for experiments 11-14 days after AAV-FLEX-TRAP virus injection.

### Translating ribosome affinity purification (TRAP)

#### Anti-GFP magnetic bead preparation

Each TRAP experiment was performed on 6 samples in parallel, with dSPN-Tsc1-WT and Tsc1-KO mice processed together. For 6 mice, two batches of beads were prepped in parallel in separate tubes. TRAP was performed according to published methods (Heiman et al., 2014; Heiman et al., 2008). All steps were performed on ice unless otherwise noted. 450 μL of Dynabeads MyOne Streptavidin T1 (Lifetech cat #65601) were washed using a magnetic tube rack in RNase-free PBS and then incubated in Protein L solution (850 μL PBS + 150 μg Protein L) for 35 min at room temperature (RT). Beads were washed 5x with 3% IgG Protease-free BSA to block, then incubated with 150 μg of two different anti-GFP antibodies (19C8 and 19F7, Memorial Sloan Kettering Antibody and Bioresource Core), diluted in 900 μL PBS, for 1 hr at RT. Beads were then washed 3x in 0.15 M KCl buffer without cyclohexamide (−CHX), then resuspended in 630 μL of 0.15 M KCl (+CHX).

#### Brain lysate preparation and immunoprecipitation

Mice were anesthetized, and brains were dissected and blocked to contain mainly striatum. Bilateral striata from each animal were placed into glass homogenization tubes on ice, and pestles were inserted. Homogenization took place at 4°C (3 strokes at 300 RPM, 12 strokes at 900 RPM, Yamato Lab Stirrer LT400), care was taken to avoid generating bubbles. Lysates were poured into pre-chilled Eppendorf tubes and centrifuged for 10 min at 2,000 x g at 4 °C to precipitate large organelles. Samples were then transferred to a new pre-chilled tube and volumes were measured. 10% NP-40 (1/9 sample volume, ∼70-80 μL) was added, then DHPC (1/9 new sample volume) was added, and samples were incubated on ice for 5 mins. DHPC stock is 200 mg dissolved in 1.38 mL ddH2O. Samples were then centrifuged for 10 min at 16,000 x g at 4 °C to precipitate mitochondria. Antibody-bound beads were resuspended by inversion, and 200 μL of beads were added to 6 separate tubes. Supernatants from samples were then transferred into tubes with beads and incubated on rotators at 4 °C overnight.

#### Isolation and purification of RNA

Samples were spun down briefly and placed on magnets pre-chilled on ice. Supernatants were collected and transferred to pre-chilled “unbound” sample tubes. Beads were washed 4x with 0.35 M KCl buffer, with samples sitting on ice for 1 min between washes to reduce background binding. Beads were resuspended in 350 μL RLT-beta-ME. 100 μL of unbound samples were added separately to 350 μL RLT-beta-ME. Samples (bound and unbound) were then rotated for 10 min at RT. Samples were placed on the magnet and supernatants were removed and added into fresh tubes containing 350 μL of 80% EtOH, mixed, and then all 700 μL of sample + EtOH was added to an RNeasy column (pre-chilled). 350 μL of unbound sample was also mixed with 350 μL of EtOH and added to RNeasy column. At this point there were 12 columns, one bound and one unbound sample for each mouse. Samples were centrifuged for 30 sec at 8000 x g at RT. Flow-through was passed through the column twice more to repeat binding. Flow-through was then discarded and 500 μL of RPE buffer was added to each column and spun for 30 sec at 8000 x g. Flow-through was discarded and 500 μL of 80% EtOH was added to the column, spun for 2 min at 8000 x g at RT. Flow-through was discarded, and columns were dried by spinning for 5 mins at full speed with cap open. Dried columns were placed into new collection tubes (not pre-chilled) and 28 μL RNase-free water was added directly to the column membrane. Columns were incubated for 5 min at RT with caps closed, then spun for 1 min at max speed at RT. RNA concentration and quality was determined by NanoDrop and Bioanalyzer at the UC Berkeley Functional Genomics Laboratory core facility.

### Quantitative PCR

Reverse transcription was performed using random hexamer primers and Superscript III reverse transcriptase (Lifetech, 18080051). Real-time PCR was performed in triplicate with 1 uL cDNA using a Biorad CFX384 thermal cycler with TaqMan Universal PCR Master Mix, no AmpErase UNG (Lifetech, 4324018) and TaqMan probes. The following TaqMan probes were used: *Grm1* (Mm01187086_m1), *Grm5* (Mm00690332_m1), *Homer1* (Mm00516275), *Plcb1* (Mm00479998), *Drd1* (Mm01353211_m1), *Drd2* (Mm00438545_m1), and *Actb* (Mm02619580_g1). Values for all mRNAs were normalized to *Actb* for each sample.

### Experimental design and statistical analyses

Experiments were designed to compare the main effect of genotype within each mouse line. The sample sizes were based on prior studies and are indicated in the figure legend for each experiment. GraphPad Prism version 8.1.2 was used to perform statistical analyses. Two-tailed paired or unpaired t-tests were used for comparisons between two groups, and an F-test was used to confirm that variances were not significantly different. For data that did not pass the D’Agostino & Pearson normality test, a Mann Whitney test was used. A one-way ANOVA with Holm-Sidak’s post-hoc tests was used to compare the means of three or more groups. For data that did not pass the D’Agostino and Pearson normality test, a Kruskal-Wallis test with Dunn’s post-hoc tests was used. A two-way ANOVA or mixed effects analysis was used to compare differences between groups for experiments with two independent variables. P-values were corrected for multiple comparisons.

## Supplemental Information - Figures and Legends

**Figure S1.**
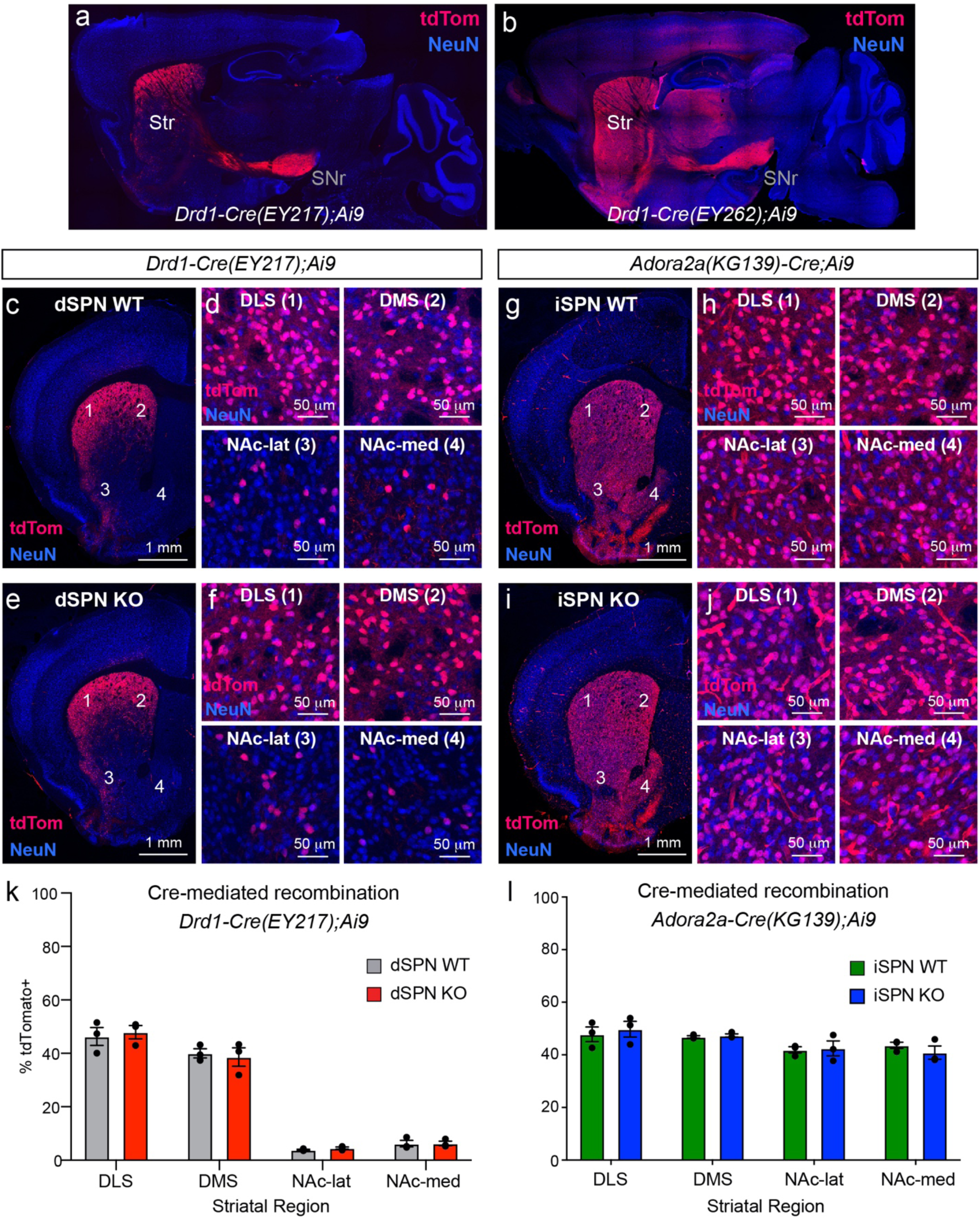
Cre-mediated recombination in *Drd1-Cre* and *Adora2a-Cre* mice. (a,b) Images of sagittal brain sections showing Cre-dependent tdTomato expression in a *Drd1-Cre(EY217);Ai9* mouse (a) and a *Drd1-Cre(EY262);Ai9* mouse (b), note striatal-restricted expression in the EY217 line. (c-f) Confocal images of coronal brain sections showing Cre-dependent tdTomato expression in a *Tsc1^wt/wt^;Drd1-Cre;Ai9* mouse (c,d) and a *Tsc1^fl/fl^;Drd1-Cre;Ai9* mouse (e,f). Higher magnification images (d,f) show region-specific expression patterns in the dorsolateral striatum (DLS, site #1), dorsomedial striatum (DMS, site #2), nucleus accumbens lateral shell (NAc-lat, site #3), and nucleus accumbens medial shell (NAc-med, site #4). (g-j) Confocal images of coronal brain sections showing Cre-dependent tdTomato expression in a *Tsc1^wt/wt^;A2a-Cre;Ai9* mouse (g,h) and a *Tsc1^fl/fl^;A2a-Cre;Ai9* mouse (i,j). Higher magnification images (h,j) show region-specific expression patterns. (k,l) Quantification of tdTomato regional expression patterns in *Tsc1^wt/wt^;Drd1-Cre;Ai9* and *Tsc1^fl/fl^;Drd1-Cre;Ai9* mice (k) and in *Tsc1^wt/wt^;A2a-Cre;Ai9* and *Tsc1^fl/fl^;A2a-Cre;Ai9* mice (l). Bars represent mean +/− SEM and dots represent individual mice. Data are expressed as the percentage of NeuN+ cells that are tdTomato+ in a given region. n=3 mice per genotype.

**Figure S2.**
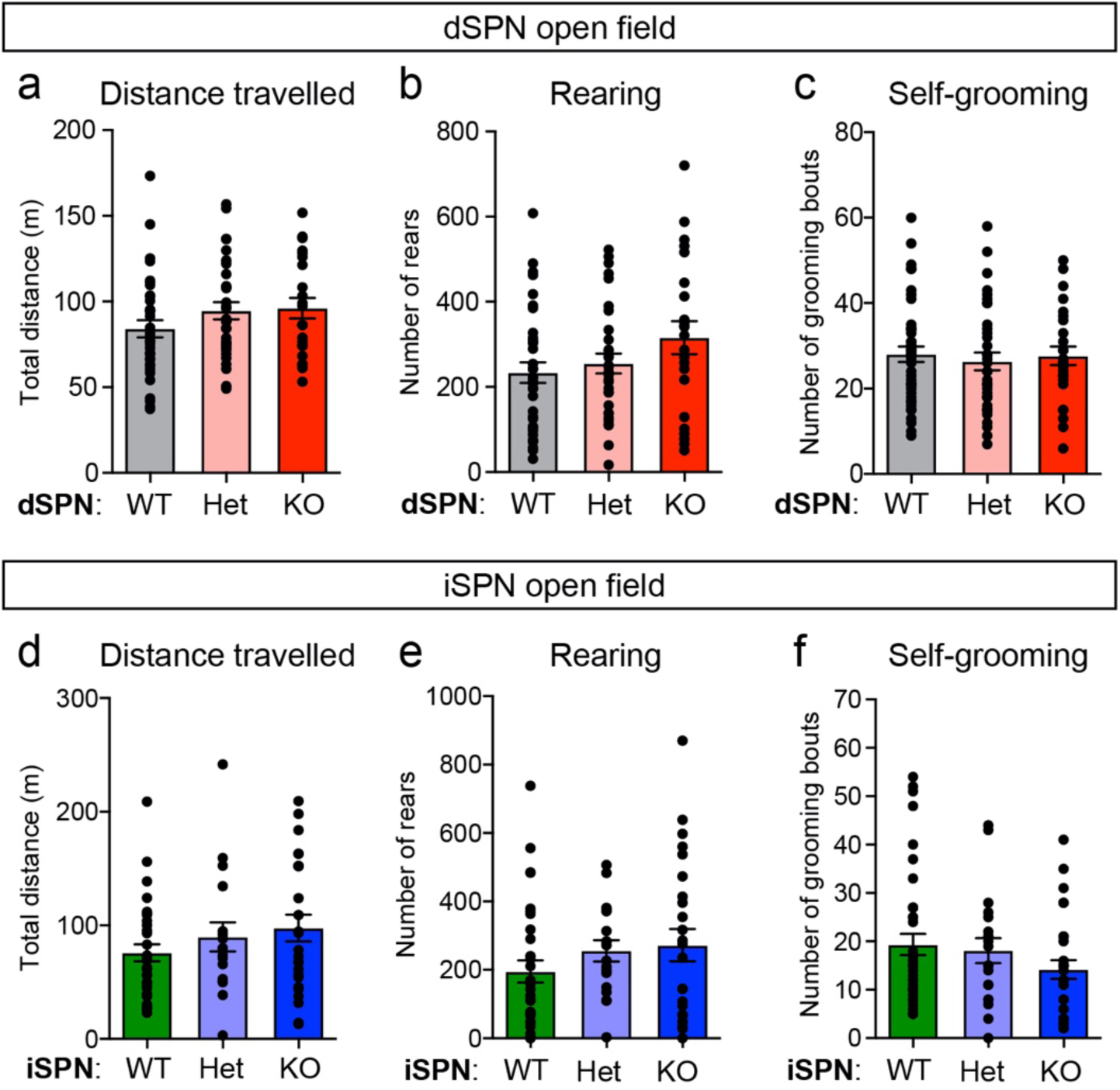
Loss of Tsc1 from SPNs does not alter open field behavior. (a-f) Quantification of open field parameters in dSPN- (a-c) and iSPN- (d-f) Tsc1 WT, Het, and KO mice. Distance travelled is the total distance travelled in the open field over 60 minutes. Rearing is the number of rears in 60 minutes. Self-grooming is the number of grooming bouts in the first 20 minutes of the open field test. Bars represent mean +/− SEM. Dots represent individual mice. dSPN-WT n=36, dSPN-Het n=32, dSPN-KO n=23, iSPN-WT n=32, iSPN-Het n=18, iSPN-KO n=25 mice. For panel a, p=0.2006; panel b, p=0.1252 (one-way ANOVA); panel c, p=0.8112 (one-way ANOVA); panel d, p=0.4028; panel e, p=0.2367; panel f, p=0.2497; Kruskal-Wallis tests except where noted.

**Figure S3.**
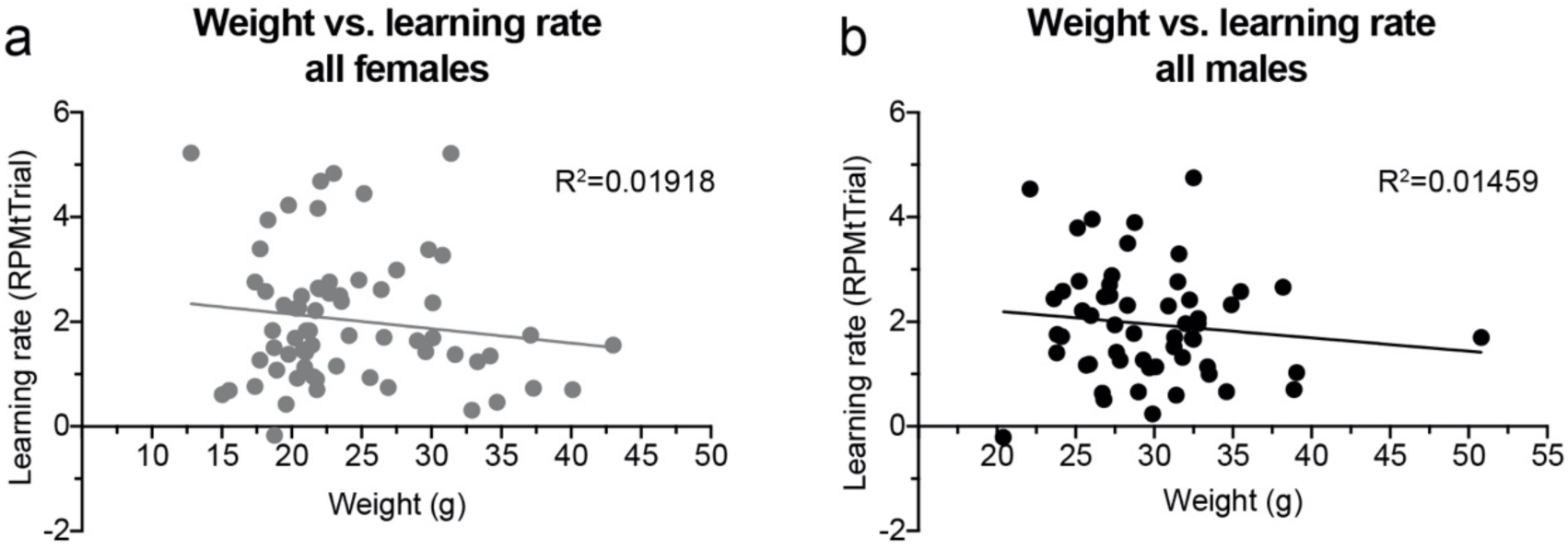
Weight does not strongly impact learning rate on the accelerating rotarod test. (a,b) Graphs show the relationship between weight in grams (g) and overall learning rate on the rotarod test for individual female (a) and male (b) mice. Learning rate was calculated as the slope of the line of performance from trial 1 to trial 12. There was not a significant relationship between weight and rotarod learning rate for females (p=0.2674, n=66 mice) or males (p=0.3796, n=55 mice). Mice from all strains (dSPN-Tsc1, iSPN-Tsc1, and Tsc2) and genotypes were combined for this analysis.

**Figure S4.**
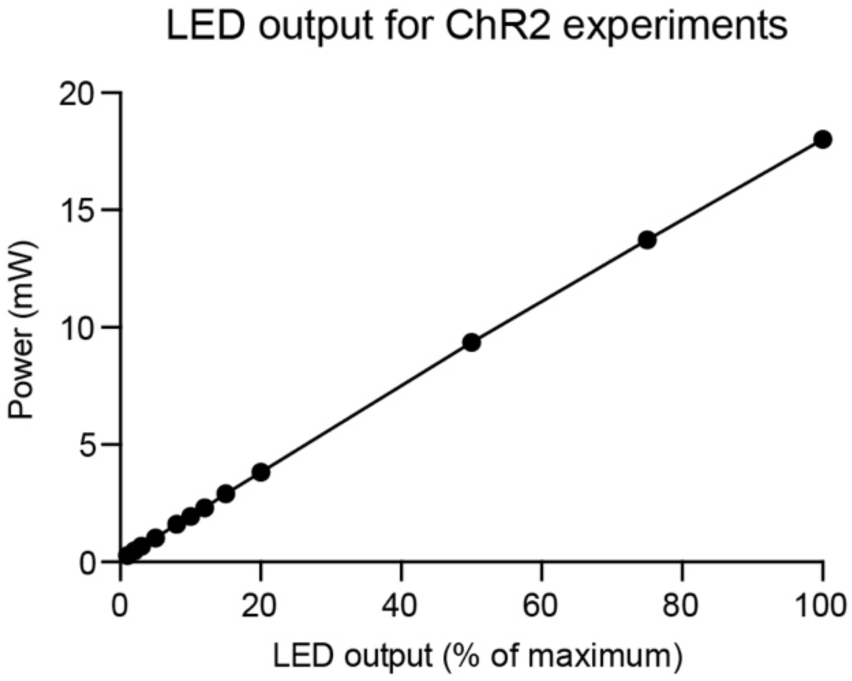
Light power plotted as a function of percent maximum LED output. 470 nm light power measured under the microscope objective shows a linear relationship with LED output.

**Figure S5.**
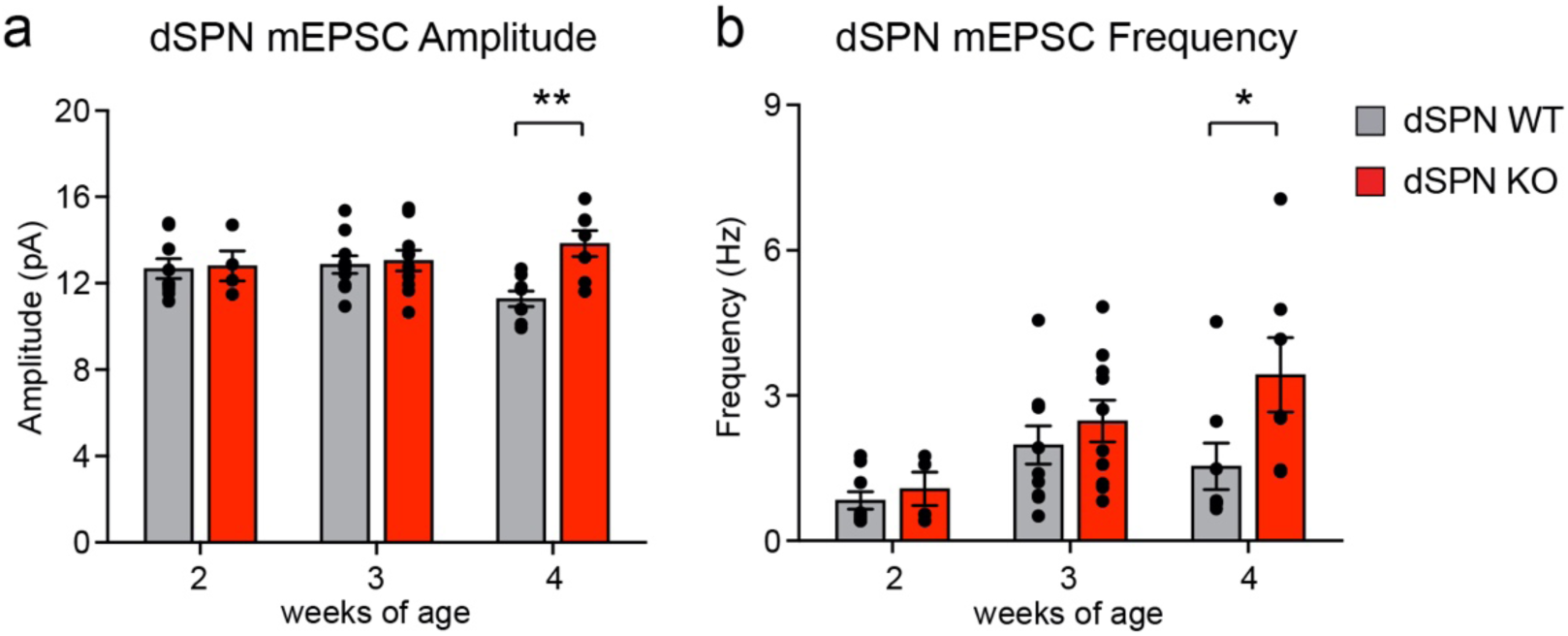
mEPSCs in dSPN-Tsc1 WT and KO mice across early postnatal development. (a,b) Mean +/− SEM mEPSC amplitude (a) and frequency (b) recorded from dSPN-Tsc1 WT and KO neurons at 2, 3 and 4 weeks of age. Dots represent individual neurons. Comparisons were made between dSPN-WT and KO at each developmental age. For panel a, p=0.8252, Mann-Whitney (2 weeks), p=0.7741, unpaired t-test (3 weeks), **, p=0.0093, Mann-Whitney (4 weeks). For panel b, p=0.7105, Mann-Whitney (2 weeks), p=0.4045, unpaired t-test (3 weeks), *, p=0.0289, Mann-Whitney (4 weeks).

**Figure S6.**
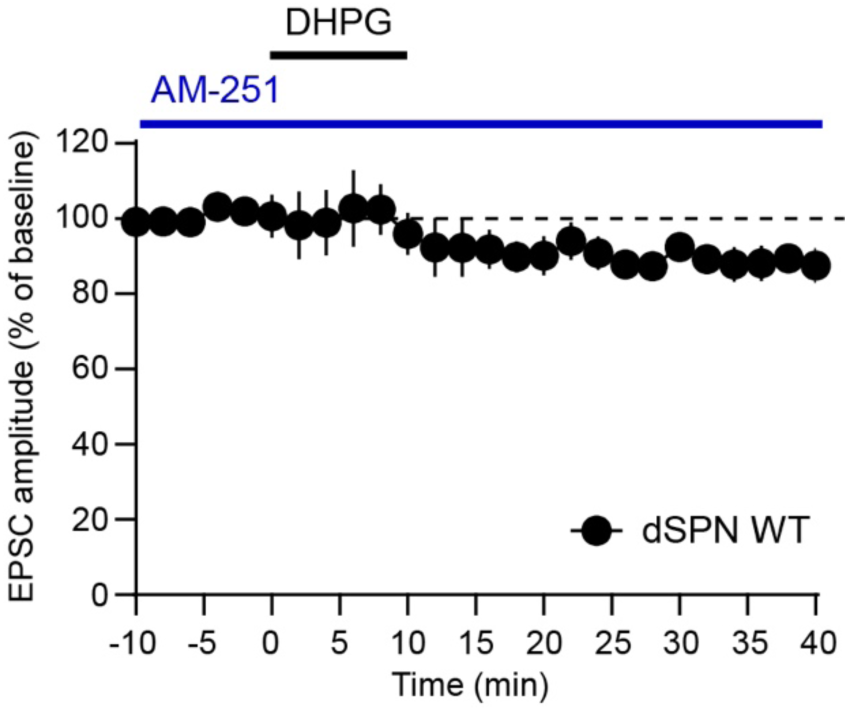
DHPG-induced LTD in WT dSPNs is blocked by a CB1 receptor antagonist. Corticostriatal terminals were stimulated with blue light (30s ISI) to evoke EPSCs in dSPN-Tsc1 WT neurons (from *Tsc1^wt/wt^;Thy1-ChR2-EYFP*;*Drd1-Cre;Ai9* mice). Following a 10-minute baseline period, DHPG (100 μM) was washed on for 10 minutes in the continued presence of AM-251 (10 μM) to block presynaptic CB1Rs. Data are presented as mean +/− SEM percent of baseline. n=5 neurons.

**Supplemental Table 1.** Summary of behavior data by sex and genotype.

| **Group/Mesaure** | **N**<br>**(mice)** | **Mean** | **Std. Deviation** | **Std. Error of**<br>**Mean** |
| --- | --- | --- | --- | --- |
| ***Open Field Test*** |  |  |  |  |
| dSPN WT Male Grooming | 16 | 26.94 | 11.91 | 2.977 |
| dSPN WT Female Grooming | 20 | 24.10 | 11.61 | 2.596 |
| dSPN Het Male Grooming | 13 | 21.00 | 12.98 | 3.600 |
| dSPN Het Female Grooming | 19 | 26.95 | 9.902 | 2.272 |
| dSPN Mut Male Grooming | 10 | 20.10 | 9.723 | 3.075 |
| dSPN Mut Female Grooming | 13 | 29.46 | 7.253 | 2.012 |
| dSPN WT Male Distance | 16 | 95.32 | 28.42 | 7.105 |
| dSPN WT Female Distance | 20 | 72.11 | 30.76 | 6.878 |
| dSPN Het Male Distance | 13 | 101.0 | 33.71 | 9.348 |
| dSPN Het Female Distance | 19 | 90.39 | 24.17 | 5.544 |
| dSPN Mut Male Distance | 10 | 89.42 | 24.47 | 7.739 |
| dSPN Mut Female Distance | 13 | 101.4 | 31.81 | 8.821 |
| dSPN WT Male Rearing | 16 | 303.8 | 156.2 | 39.05 |
| dSPN WT Female Rearing | 20 | 171.3 | 112.7 | 25.19 |
| dSPN Het Male Rearing | 13 | 295.2 | 163.7 | 45.40 |
| dSPN Het Female Rearing | 19 | 226.9 | 101.5 | 23.30 |
| dSPN Mut Male Rearing | 10 | 288.6 | 163.0 | 51.53 |
| dSPN Mut Female Rearing | 13 | 336.6 | 203.6 | 56.48 |
| iSPN WT Male Grooming | 20 | 30.30 | 30.60 | 6.842 |
| iSPN WT Female Grooming | 24 | 31.22 | 31.71 | 6.473 |
| iSPN Het Male Grooming | 11 | 20.09 | 10.89 | 3.282 |
| iSPN Het Female Grooming | 7 | 10.86 | 7.010 | 2.650 |
| iSPN Mut Male Grooming | 14 | 35.37 | 35.52 | 9.494 |
| iSPN Mut Female Grooming | 11 | 17.60 | 11.16 | 3.366 |
| iSPN WT Male Distance | 20 | 78.92 | 39.74 | 8.887 |
| iSPN WT Female Distance | 24 | 74.93 | 47.13 | 9.621 |
| iSPN Het Male Distance | 11 | 89.94 | 56.66 | 17.08 |
| iSPN Het Female Distance | 7 | 89.76 | 54.43 | 20.57 |
| iSPN Mut Male Distance | 14 | 94.99 | 54.82 | 14.65 |
| iSPN Mut Female Distance | 11 | 100.8 | 66.66 | 20.10 |
| iSPN WT Male Rearing | 20 | 218.2 | 140.2 | 31.34 |
| iSPN WT Female Rearing | 24 | 192.3 | 199.2 | 40.66 |
| iSPN Het Male Rearing | 11 | 223.4 | 100.1 | 30.19 |
| iSPN Het Female Rearing | 7 | 306.7 | 171.1 | 64.67 |
| iSPN Mut Male Rearing | 14 | 314.7 | 249.6 | 66.71 |
| iSPN Mut Female Rearing | 11 | 218.2 | 219.1 | 66.06 |
| ***Rotarod*** |  |  |  |  |
| dSPN WT Male Learning Rate | 15 | 1.462 | 0.9233 | 0.2384 |
| dSPN WT Female Learning Rate | 16 | 1.096 | 1.009 | 0.2522 |
| dSPN Het Male Learning Rate | 10 | 1.398 | 0.9793 | 0.3097 |
| dSPN Het Female Learning Rate | 15 | 2.171 | 1.064 | 0.2746 |
| dSPN KO Male Learning Rate | 19 | 1.967 | 1.218 | 0.2794 |
| dSPN KO Female Learning Rate | 14 | 2.421 | 1.294 | 0.3458 |
| iSPN WT Male Learning Rate | 17 | 1.320 | 1.004 | 0.2435 |
| iSPN WT Female Learning Rate | 17 | 1.430 | 1.023 | 0.2482 |
| iSPN Het Male Learning Rate | 7 | 1.513 | 0.6812 | 0.2575 |
| iSPN Het Female Learning Rate | 6 | 1.402 | 1.168 | 0.4766 |
| iSPN KO Male Learning Rate | 9 | 1.857 | 0.8035 | 0.2678 |
| iSPN KO Female Learning Rate | 9 | 1.295 | 0.8304 | 0.2768 |
| Tsc2 WT Male Learning Rate | 5 | 1.742 | 1.432 | 0.6405 |
| Tsc2 WT Female Learning Rate | 12 | 1.388 | 0.8415 | 0.2429 |
| Tsc2 Het Male Learning Rate | 12 | 2.604 | 1.043 | 0.3012 |
| Tsc2 Het Female Learning Rate | 19 | 2.534 | 1.524 | 0.3497 |

